# Functional Connectome of Superagers Reveals Early Markers of Resilience and Vulnerability to Alzheimer’s Disease

**DOI:** 10.1101/2025.07.20.665707

**Authors:** Kanhao Zhao, Hua Xie, Gregory A. Fonzo, Nancy B. Carlisle, Tovia Jacobs, Ricardo S. Osorio, Arpana Church, Feng Vankee Lin, Yu Zhang, the ADNI Study Group

**Author notes:** Corresponding author: Yu Zhang, Assistant Professor of Psychiatry and Behavioral Sciences, Stanford University School of Medicine, Stanford, CA 94305, USA. Data used in preparation of this article were obtained from the Alzheimer’s Disease Neuroimaging Initiative (ADNI) database (adni.loni.usc.edu). As such, the investigators within the ADNI contributed to the design and implementation of ADNI and/or provided data but did not participate in analysis or writing of this report. A complete listing of ADNI investigators can be found at: http://adni.loni.usc.edu/wp-content/uploads/how_to_apply/ADNI_Acknowledgement_List.pdf.

## Abstract

As populations age, identifying the neurobiological basis of cognitive resilience is critical for delaying or preventing Alzheimer’s disease (AD). While most older adults experience memory decline, a subset known as superagers (SA) maintains youthful memory into late life, offering a unique window into protective mechanisms against neurodegeneration. Here, we identified a functional connectivity (FC) signature, termed Alzheimer’s-resilient connectome (ARC), that robustly differentiates SA from age-matched patients with AD. Using resting-state fMRI in a discovery cohort (N = 290), we identified ARC derived from machine learning classifiers that distinguished SA from AD with high accuracy (AUC = 0.85), and validated the replicability of the ARC in an independent replication cohort (N = 143). ARC involved prefrontal, temporal and insular networks and was strongly associated with brain age. When applied to cognitively unimpaired (CU) adults (discovery cohort: N = 818 and replication cohort: N = 497), ARC-based subtyping revealed SA-like and AD-like subgroups with similar baseline cognitive performance but markedly divergent longitudinal trajectories. SA-like CU individuals showed slower cognitive decline, reduced amyloid-β accumulation, and lower risk of conversion to mild cognitive impairment and AD, reinforcing the ARC signature as a potential early indicator of resilience. Genome-wide association analysis identified *CLYBL* and *FRMD6* as novel genetic modulators associated with these divergent aging phenotypes. Together, our findings position ARC as a sensitive and generalizable biomarker of resilience, enabling early risk stratification and precision prevention for AD.

## Introduction

Brain aging is a complex biological process influenced by various factors, including chronological age, genetic risk, and dynamic changes in brain function^1,2^. As individuals age, cognitive decline, particularly in episodic memory, is commonly observed, especially among those with Alzheimer’s Disease (AD) related dementia^3,4^. Identifying mechanisms that can slow or prevent memory decline is a top priority in the field of AD research.

To date, prevention efforts have primarily focused on individuals with mild cognitive impairment (MCI), a critical transitional stage preceding dementia^5,6^. However, the symptomatic manifestation at the MCI stage may indicate that it is already late for effective intervention. In contrast, CU individuals, although symptom-free, may harbor underlying AD neuropathological changes such as amyloid-β (Aβ) accumulation^8,9^. Compared to CU without AD neuropathological changes, these individuals show a high risk of progressing to AD, making this stage a critical window for preventive efforts. Notably, emerging literature suggests that a subset of older individuals appear to resist such pathological changes, maintaining high cognitive function and showing neurobiological characteristics akin to those of a distinct group known as superagers (SA), who retain memory performance comparable to middle-aged or young-old groups^9–11^. This cognitive resilience, often referred to as cognitive reserve, may reflect underlying protective mechanisms that delay or prevent progression to AD^10^. Understanding the mechanistic differences that explain memory retention in SA compared to those with AD-related dementia may provide insights into brain aging in general^12,13^. Specifically, for CU individuals, this knowledge could help clarify who is at risk of developing AD-related dementia.

Functional connectivity (FC), measuring the temporal correlation in functional magnetic resonance imaging (fMRI) time series between brain regions, offers insights into large-scale brain network architecture and has emerged as a promising modality for studying the neural mechanisms underlying cognitive aging^14^. Prior studies have demonstrated that specific FC patterns are associated with cognitive resilience and memory performance in older adults^15,16^. For instance, SA shows robustly connected FC within the default mode network (DMN), a network implicated in episodic memory and typically disrupted in aging and AD^17,18^. Other studies using FCs to chart brain aging have found its association with cognitive performance and AD pathology in CU individuals^19^, highlighting the potential of FC as a biomarker of latent risk or resilience. This suggests that studying FC profiles in SA and identifying similar patterns in CU may offer a unique opportunity to identify connectome-based protective signatures that can be used for early detection of individuals who are vulnerable or resilient to AD pathology and which may serve as therapeutic targets for preventing the disease. However, prior studies have primarily focused on known networks or selected contrasts and have not systematically examined whether FC signatures can be used to predict aging trajectories and stratify CU individuals in a clinically meaningful and generalizable manner.

In parallel, recent advances in genetic studies have deepened our understanding of cognitive decline and resilience. Genome-wide association studies (GWAS) have identified risk loci, such as APOE ε4 and *BIN1*, which implicate synaptic, inflammatory, and lipid metabolic pathways in AD pathogenesis^20,21^. Notably, some variants appear to support cognitive resilience independent of amyloid or tau burden, suggesting distinct protective mechanisms^22–24^. These findings highlight the value of polygenic approaches for understanding individual variability in brain aging. Yet, few studies have associated genetic variation with FC-based biomarkers to clarify mechanisms that distinguish resilient from vulnerable trajectories in CU individuals. Addressing this gap could offer novel insights into the molecular architecture of functional brain aging and resilience.

Building on this background, we hypothesized that the FC patterns differentiating SA from individuals with AD reflect underlying mechanisms of brain aging and could be used to stratify CU individuals into biologically distinct subgroups with varying risks of cognitive decline and neuropathological progression. To test this hypothesis, we constructed a discovery cohort (N = 290; 180 SA and 110 age-matched AD patients) using data from the OASIS-3 and HABS datasets, and used the ADNI dataset as an independent replication cohort (N = 143; 64 SA and 79 age-matched AD patients). We then trained sparse support vector machine (SVM) classifiers on resting-state FC data from the discovery cohort to identify a FC signature, termed Alzheimer’s-resilient connectome (ARC), that robustly differentiates SA from AD. The ARC signature demonstrates a strong correlation with brain aging and various cognitive and emotion regulation functions. The accuracy of the signature was rigorously validated through cross-validation and independently confirmed in the replication cohort. We then applied the ARC signature to stratify CU individuals from both the discovery (N = 818) and replication (N = 497) cohorts into SA-like and AD-like subtypes. Although these subtypes show comparable baseline cognitive performance, they exhibit marked differences in longitudinal outcomes. Compared to SA-like CU individuals, AD-like ones experienced faster memory decline, increased Aβ accumulation, and higher risk of progression to MCI or AD. These subtypes also differ in brain aging trajectories and show high replicability in replication cohort. Finally, a genome-wide association study identified *CLYBL* and *FRMD6* as potential genetic modulators of the observed phenotypic divergence between subtypes, offering insights into the molecular basis of resilience and vulnerability in aging.

## Results

### ARC distinguishes superagers from individuals with AD

To investigate FC features associated with memory resilience versus neurodegeneration, we first aimed to identify robust and replicable patterns that distinguish SA from individuals with AD. This classification serves as the foundation for extracting connectomic markers of brain resilience. In our study, SA was defined based on three validated criteria^10,25^: (1) Age threshold: individuals aged ≥ 75 years; (2) Superior episodic memory performance, exceeding that of middle-aged normative adults, assessed using the *Wechsler Memory Scale (WAIS-R) logic memory IA* ≥ 13 and *IIA* ≥ 11^26,27^ in the discovery cohort, and the *Rey Auditory Verbal Learning Test (RAVLT) delay recall* ≥ 9^10,28^ in the replication cohort; and (3) Comparable executive function to age-matched older adults, determined by a *Trail Making Test Parts B* (TMT-B) score ≤ 166^29^. Details on the inclusion flow are illustrated in Figure S1 and the *Superager definition section* of the Methods. Demographics for SA and age-matched AD subjects in the discovery and replication cohorts are summarized in Tables S1 and S2.

We first calculated regional pairwise FC from resting-state fMRI time series based on the Schaefer atlas^30^ of 100 parcels (Figure S2A). Sparse support vector machine (SVM) models (see Method: *Classification between SA and AD* section) were then trained with FC features to classify SA and age-matched AD. Model evaluation using ten-fold cross-validation in the discovery cohort comprising 180 SA and 110 AD subjects demonstrates accurate classification (Figure 1A, accuracy = 0.80 ± 0.05, sensitivity = 0.74 ± 0.14, specificity = 0.83 ± 0.08, area under the receiver operating characteristic curve (AUC) = 0.85 ± 0.08). To visualize the most discriminative FCs, we averaged the SVM model weights across cross-validation folds and mapped the sum of the absolute values of weights as node strengths onto the brain surface (Figure 1B). The classifier-identified discriminative FCs involved the bilateral caudal middle frontal cortex, angular cortex, right posterior cingulate cortex, bilateral precuneus from DMN, postcentral cortex, insula from sensorimotor network (SMN), and left anterior cingulate cortex from ventral attention network (VAN). To test the insensitivity of classification results to potential site effects, we separately evaluated model performance on OASIS-3 and HABS datasets of the discovery cohort, yielding accuracies of 0.78 ± 0.09 and 0.82 ± 0.09 (Figure S3A). Furthermore, we tested the generalizability of the trained models on an independent replication cohort (N = 143, 64 SA and 79 AD). Predicted labels, derived from averaged classifier confidence scores, yielded an accuracy of 0.71, sensitivity of 0.71, specificity of 0.71, and AUC of 0.72 (Figure 1A), confirming robust cross-cohort performance.

**Figure 1.**
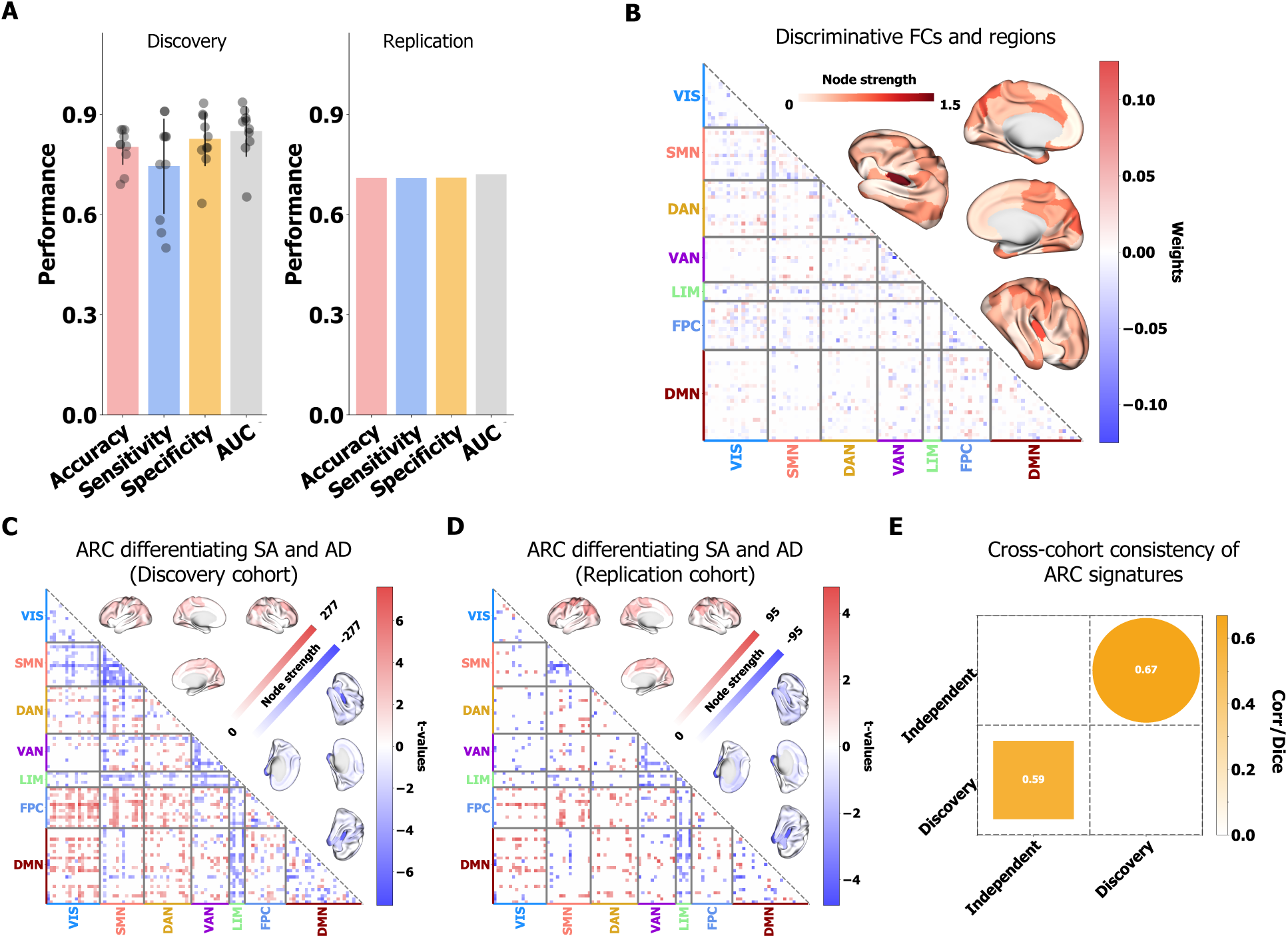
Identification and validation of the Alzheimer’s-resilient connectome (ARC) distinguishing superagers from individuals with AD. **(A)** Classification performance in the discovery and replication cohorts. Ten-fold cross-validation in the discovery cohort yielded robust accuracy = 0.80 ± 0.05, sensitivity = 0.74 ± 0.14, specificity = 0.83 ± 0.08, and area under the curve (AUC) = 0.85 ± 0.08. The trained models were then applied to the replication cohort, where predicted labels derived from averaged confidence scores achieved accuracy = 0.71, sensitivity = 0.71, specificity = 0.71, and AUC = 0.72. **(B)** Discriminative FC signature based on the average weights of the ten cross-validated classifiers. Node strength, defined as the sum of absolute FC classifier weights for each region, is visualized on the brain surface to highlight regions contributing most to classification. **(C), (D)** ARC signatures that significantly differentiated SA from AD as detected by two-sample t-tests (FDR-corrected p < 0.05) in the discovery and replication cohorts. Hypo-connections (stronger in SA) are colored red, while hyper-connections (stronger in AD) are colored blue. Node strength, derived from summed positive and negative t-values of distinguishing FCs, is visualized on the brain surface. **(E)** Cross-cohort consistency of ARC signatures. Heatmaps display Kendall’s rank correlation and Dice coefficient. The upper triangular heatmap represents Kendall correlation coefficients (circle box), while the lower triangular heatmap represents Dice coefficients (rectangle box). All the values in the heatmap were significant (p < 0.001).

Next, we investigated how these discriminative FCs in SA were differentiated from AD. Two-sample t-tests were conducted on the discovery cohort to compare these FCs, categorizing them into hypo-connections (stronger in SA; red in Figure 1C, FDR-corrected p < 0.05) and hyper-connections (stronger in AD; blue, FDR-corrected p < 0.05). Hypo-connections were primarily located in the bilateral caudal middle frontal cortex, angular cortex, and bilateral postcentral cortex, whereas the hyper-connections involved the right insula and bilateral temporal poles. These FC patterns collectively constituted the Alzheimer’s-resilient connectome (ARC) signature. To confirm the robustness of ARC, we applied the same statistical comparison within the OASIS-3 and HABS subsets to obtain ARC signature of each dataset from discovery cohort (Figure S3B, C) and then quantified the consistency of these signatures (Figure S3D), using Kendall rank correlation and Dice coefficient, with statistical significance evaluated via 1,000 permutation tests. We next used the same statistical measurements to identify the ARC signature in the replication cohort (Figure 1D) and quantified the reproducibility of ARC signature between discovery and replication cohorts (Figure 1E). Overall, the identified ARC was highly consistent across all datasets and cohorts (τ = 0.67, p < 0.001, Dice Index = 0.59), supporting the stability and generalizability of ARC as a distinctive connectomic signature of resilience.

### Phenotypic profiles associated with ARC scores

To further characterize the functional relevance of the ARC and gain a comprehensive understanding of the cognitive domains linked to ARC in SA, we examined its associations with a wide range of phenotypic variables. These measures included demographic variables, APOE genotype, and cognitive domains assessed through the *Clinical Dementia Rating Scale* (CDR)^31^, *Functional Activities Questionnaire* (FAQ)^32^, *Neuropsychiatric Inventory* (NPS)^33^, and a neuropsychological assessment battery (NAB)^34^. For continuous variables, we computed Kendall rank correlations with the ARC scores derived from ARC-signature-weighted sum of FCs, where the age and sex were set as covariates. For binary categorical variables, we used the Kruskal–Wallis test. All p-values were corrected for multiple comparisons using the false discovery rate (FDR), separately for the discovery and replication cohorts. As shown in Figure 2 (A, C), the ARC scores were significantly associated with age and sex. In addition to expected associations with memory-related domains—which were part of the SA definition—the ARC was also linked to multiple cognitive and functional domains beyond episodic memory. Specifically, ARC scores correlated with CDR subdomains such as judgment and community affairs, FAQ items including tax affairs, traveling, and driving, and performance of neuro-cognitive tests like *Boston Naming Test* (BNT) (Figure 2 (B, D))^35^. Additionally, significant correlations were also observed between the ACR scores and NPS including depression and apathy. Importantly, these associations observed in the discovery cohort were consistently reproducible in the replication cohort.

**Figure 2.**
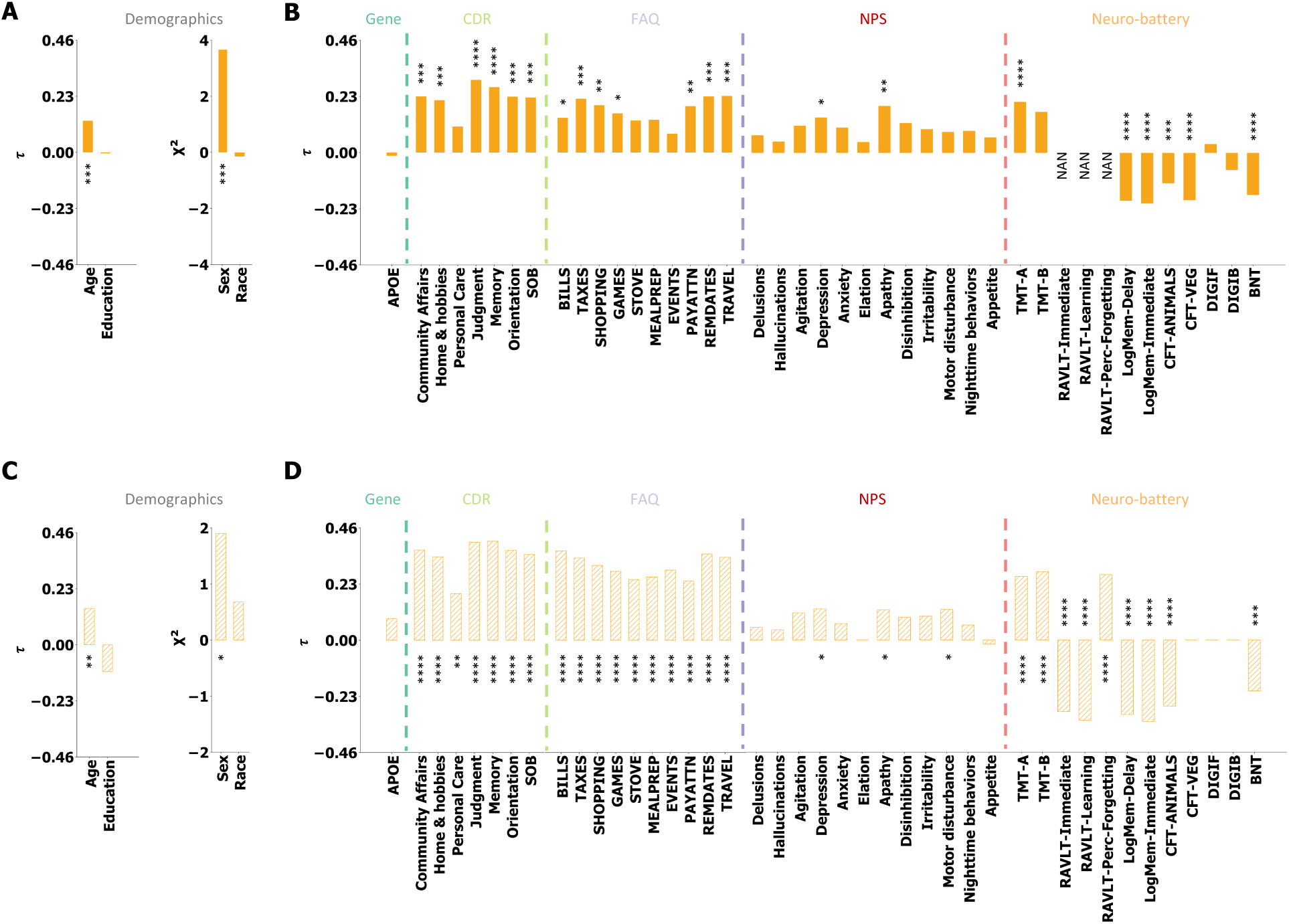
Phenotypic profiles associated with ARC. The association between various phenotypic indices and the weighted sum of the ARC was examined across SA and AD, forming a comprehensive phenotypic profile. Kendall rank correlation was used to assess relationships with continuous variables, while the Kruskal–Wallis test was applied to categorical variables. For non-demographic variables, Kendall rank correlation analyses were adjusted for age and sex as covariates. FDR correction was applied to all p-values from the discovery and replication cohorts separately, with significance levels denoted as * (p_fdr_ ≤ 0.05), ** (p_fdr_ ≤ 0.01), *** (p_fdr_ ≤ 0.001), and **** (p_fdr_ ≤ 0.0001). **(A)** Association with demographic variables in the discovery cohort. **(B)** Association with APOE genotype and cognitive variables in the discovery cohort. **(C)** Association with demographic variables in the replication cohort. **(D)** Association with APOE genotype and cognitive variables in the replication cohort.

Next, we examined whether the association between the ARC scores and cognitive performance generalized across the broader cognitive spectrum, including SA, CU, and AD individuals. Specifically, in the replication cohort, we applied the ARC signature to baseline CU data to compute individual ARC scores and assessed their relationship with various cognitive measures using Kendall rank correlation. Given that CU participants were, on average, younger than those in the SA and AD groups, age was included as a covariate in the partial correlation analysis to account for the potential confounding. As shown in Figure S4, the ARC scores were significantly correlated with indicators of daily functional impairment and memory decline, supporting its relevance to cognitive function across a continuum— from cognitively resilient to impaired individuals.

### ARC scores reflect individual differences in brain age

Accelerated brain aging is a well-established feature of AD^36,37^, whereas SA tends to exhibit preserved, youth-like brain characteristics despite the advanced age^38,39^. Based on the prior evidence, we hypothesized that the ARC would be closely associated with individual variation in predicted brain age. To test this, we trained a Bayesian ridge regression model^40^ to predict chronological age based on FC features in CU from the discovery cohort. Model performance was assessed through ten-fold cross-validation and further validated by applying the trained models to the replication cohort. The model exhibited robust predictive accuracy, as assessed by R² and Pearson correlation coefficients between predicted brain age and chronological age (Figure 3A; discovery cohort: R^2^ = 0.35, r = 0.60, p = 1.5 × 10^−78^; replication cohort: R^2^ = 0.24, r = 0.50, p = 1.4 × 10^−32^).

**Figure 3.**
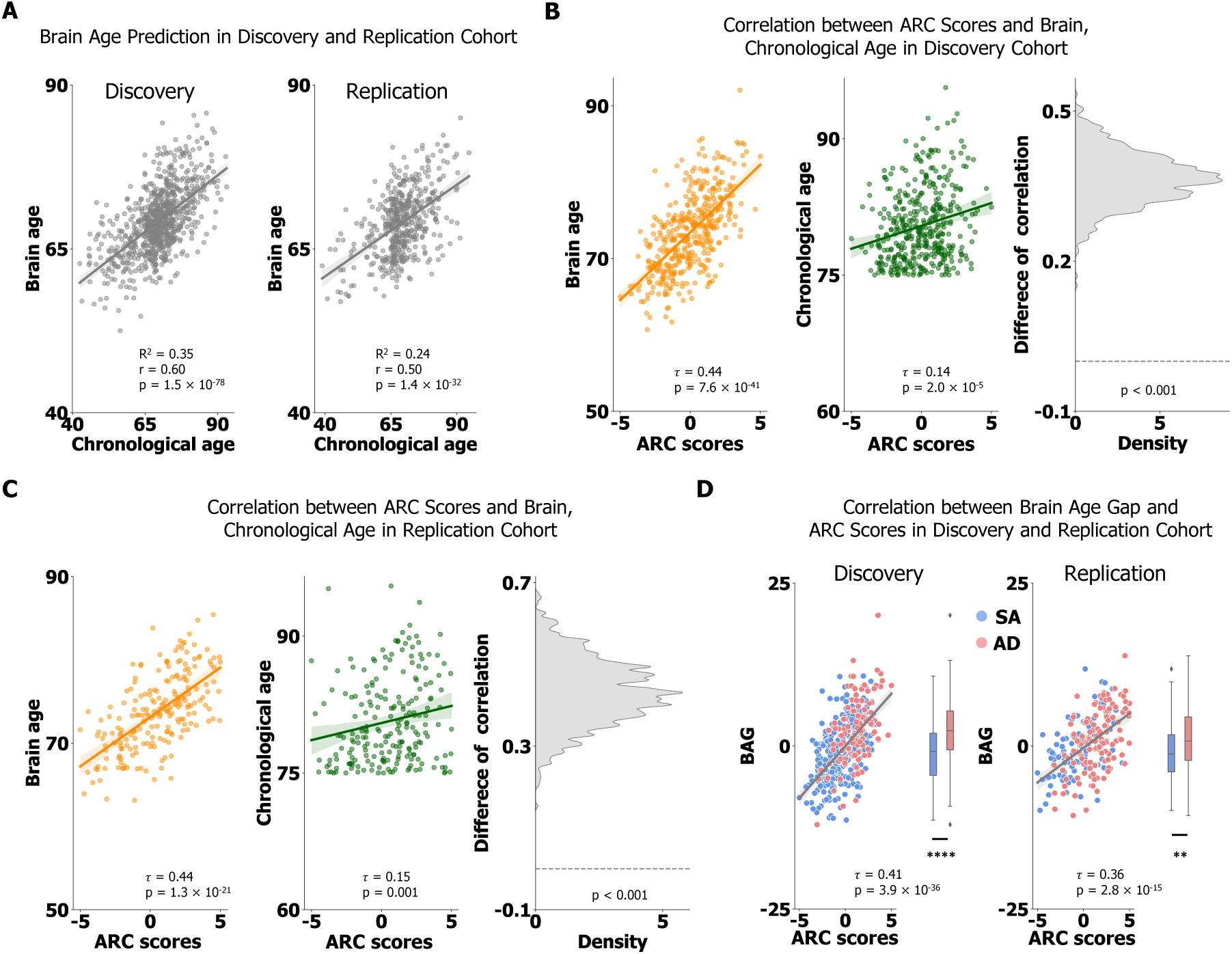
Association between the ARC scores and brain aging in SA and AD. **(A)** Brain age prediction performance assessed in the discovery and replication cohorts. A Bayesian ridge regression model was trained using whole-brain FC features from CU individuals in discovery cohort to predict chronological age. Model performance was evaluated using ten-fold cross-validation, and further tested in the replication cohort using cross-validated modals. Predictive accuracy was assessed using R² and Pearson correlation between predicted brain age and chronological age (discovery cohort: R^2^ = 0.35, r = 0.60, p = 1.5 × 10^−78^; replication cohort: R^2^ = 0.24, r = 0.50, p = 1.4 × 10^−32^). **(B), (C)** Correlations between ARC signature scores (z-scored weighted sum of ARC FCs) and predicted brain age versus chronological age across SA and AD individuals. In both the discovery and replication cohorts, predicted brain age was more strongly correlated with ARC signature scores (discovery: *τ* = 0.44 vs. 0.14; replication: τ = 0.44 vs. 0.15). To enhance statistical power, multiple sessions from the same subject were included, with each dot representing a sample rather than a subject (n = 414 for the discovery cohort, and n = 213 for the replication cohort. Statistical significance was assessed via Kendall rank correlation (used due to non-normal age distributions), and differences between correlation coefficients were confirmed by 1,000 bootstrapping iterations (p < 0.001 for both cohorts). **(D)** Association between ARC signature scores and brain age gap (BAG), calculated as the difference between predicted brain age and chronological age. ARC was significantly correlated with BAG in both the discovery cohort (*τ* = 0.41, p = 3.9 × 10^−36^) and replication cohort (*τ* = 0.36, p = 2.8 × 10^−15^). Differences in BAG between SA and AD was tested via two-sample t-tests, with significance levels indicated as ** (p ≤ 0.01), and **** (p ≤ 0.0001).

Next, we applied the brain age prediction model to SA and AD subjects in both cohorts to estimate their brain ages and examined the relationship between the ARC scores and predicted brain age. In both discovery and replication cohorts, ARC showed significantly stronger correlation with the predicted brain age than with chronological age, as confirmed by 1,000 bootstrap comparisons of correlation coefficients (Figure 3B, C, p < 0.001). To further quantify the extent of brain aging, we calculated the brain age gap (BAG) as the difference between predicted brain age and chronological age, following by a linear regression to control the bias to chronological age^37^. A larger BAG indicates accelerated brain aging, whereas a smaller BAG suggests a more preserved, youthful brain. As hypothesized, ARC was significantly correlated with BAG in both the discovery and replication cohorts (Figure 3D), supporting the notion that ARC captured meaningful inter-subject differences in brain age.

### Phenotypic profiles differentiating ARC-defined CU subtypes

Since ARC captured FC patterns associated with brain resilience and aging, we next investigated whether it could differentiate CU individuals into subtypes with distinct cognitive and clinical profiles. We applied cross-validated classification models to assign subtype labels. CU individuals at baseline were classified as SA-like if they were predicted as SA in more than half of the cross-validation folds, while the remainder were classified as AD-like CU (Figure S2B). This procedure yielded 607 SA-like and 211 AD-like individuals in the discovery cohort, and 370 SA-like and 127 AD-like individuals in the replication cohort. While the subtypes were derived from the ARC-based binary classification model, it remained critical to assess the stability and reliability of these subtype assignments across modeling variations and sampling perturbations. We employed two complementary strategies to evaluate the robustness of the classification-based subtyping. First, we repeated the classification and clustering procedures across 10 replication trials, each initialized with a different random seed. Second, we performed bootstrapped subsampling by randomly selecting 90% of participants (AD and SA) in the classification step across 10 trials and reapplying the pipeline to obtain subtypes. In both strategies, the consistency of subtype assignments was assessed using the Dice coefficient across 10 trials. As shown in Figure S5, both strategies yielded high average Dice coefficients (> 0.93), demonstrating strong reliability of the ARC-based subtyping procedure^41,42^. Additionally, although the subtyping was derived from a binary classifier, we explored whether more complex subtype structures might be supported by distribution of predicted classification scores. Specifically, we applied k-means clustering to the ARC scores of CU individuals and found that the two-subtype solution consistently produced the highest stability, supporting its selection for further analysis.

### Baseline FC differences between ARC-defined SA-like and AD-like subtypes

To examine how ARC signature differentiates the SA-like and AD-like CU subtypes at baseline, we performed two-sample t-tests on FCs within the discovery cohort, controlling for age and sex. As shown in Figure S6A, the SA-like subtype exhibited significantly stronger FCs (red) involving the bilateral angular gyri and superior frontal cortices. In contrast, the AD-like subtype showed significantly stronger FCs (blue) involving the bilateral insula, orbitofrontal cortex, and temporal lobe. These patterns were reproducible in the replication cohort (Figure S6D), as confirmed by high Kendall rank correlation and Dice coefficients between cohorts (Figure S6G). Since CU subtypes were derived using models trained to distinguish SA from AD, we expected that the FC pattern separating SA-like and AD-like CU individuals (Figure S6A and D) would resemble the ARC signature itself (Figure 1C and D). Supporting this, we observed a significant correlation (Kendall’s *τ* = 0.3, p < 0.001) between the FC pattern differentiating CU subtypes and the ARC signature, indicating shared topological features between the two.

We further compared each CU subtype to its corresponding reference group (SA or AD) at baseline. In the discovery cohort, the AD-like subtype exhibited higher FCs in the left superior medial and superior frontal cortices, and lower FCs in the bilateral superior temporal cortex and anterior insula, compared to AD (Figure S6B). These results were replicated in the replication cohort (Figure S6E), with consistent FC differences confirmed by significant Kendall rank correlation and Dice coefficient values (Figure S6H). In contrast, the comparison between the SA-like subtype and the SA group revealed limited and inconsistent FC differences across the two cohorts (Figure S6C, F), suggesting that the SA-like CU subtype may closely resemble the canonical SA profile in terms of baseline FC.

### Phenotypic and longitudinal profiles differentiating ARC-defined CU subtypes

We examined whether SA-like and AD-like subtypes exhibited distinct demographic and cognitive characteristics. Using the Kruskal-Wallis test, we compared the two subtypes across demographic variables, cognitive domains from CDR and NAB, and daily functional measures from the FAQ. Baseline comparisons in both the discovery and replication cohorts are summarized in Tables S3-S5. While the executive function difference was observed specifically in the replication cohort, age was the only variable that significantly differentiated the two subtypes across both cohorts, with SA-like individuals being younger than their AD-like counterparts (Figure S7).

Given the FC differences (Figure S6) between the ARC-defined CU subtypes, we expected that these subtypes would exhibit distinct cognitive profiles at baseline, resembling the patterns observed between SA and AD. However, baseline comparisons revealed minimal cognitive differences. This could reflect that the relatively younger age of the CU participants potentially masks early cognitive variation. Additionally, standard neuropsychological measures may lack the sensitivity to detect subtle, early-stage cognitive differences in cognitively unimpaired individuals. We reasoned that cognitive differences may emerge over time, with SA-like individuals showing more preserved cognitive trajectories compared to AD-like individuals as pathological aging progresses. To test this hypothesis, we introduced accelerated longitudinal designs by applying linear mixed-effects models to examine developmental trajectory for each characteristic variable (excluding demographics) across the two subtypes. Results revealed that AD-like CU individuals exhibited significantly faster cognitive decline than SA-like CU individuals, as evidenced by greater worsening of overall CDR scores and declines in executive function, as measured by TMT scores within the NAB cognitive tests (Figure 4). Additional domain-level analyses of CDR subitems, particularly memory and attention, further supported this trend (Figure S8). Moreover, we assessed the proportion of individuals in each subtype who progressed to MCI or AD over time, observing a significantly higher conversion rate in the AD-like subtype compared to the SA-like subtype (Figure S9, discovery cohort: OR [95% CI]: 2.19 [1.24, 3.86]; independent cohort: OR [95% CI]: 1.89 [1.05, 3.39]), highlighting the prognostic utility of ARC-based subtyping.

**Figure 4.**
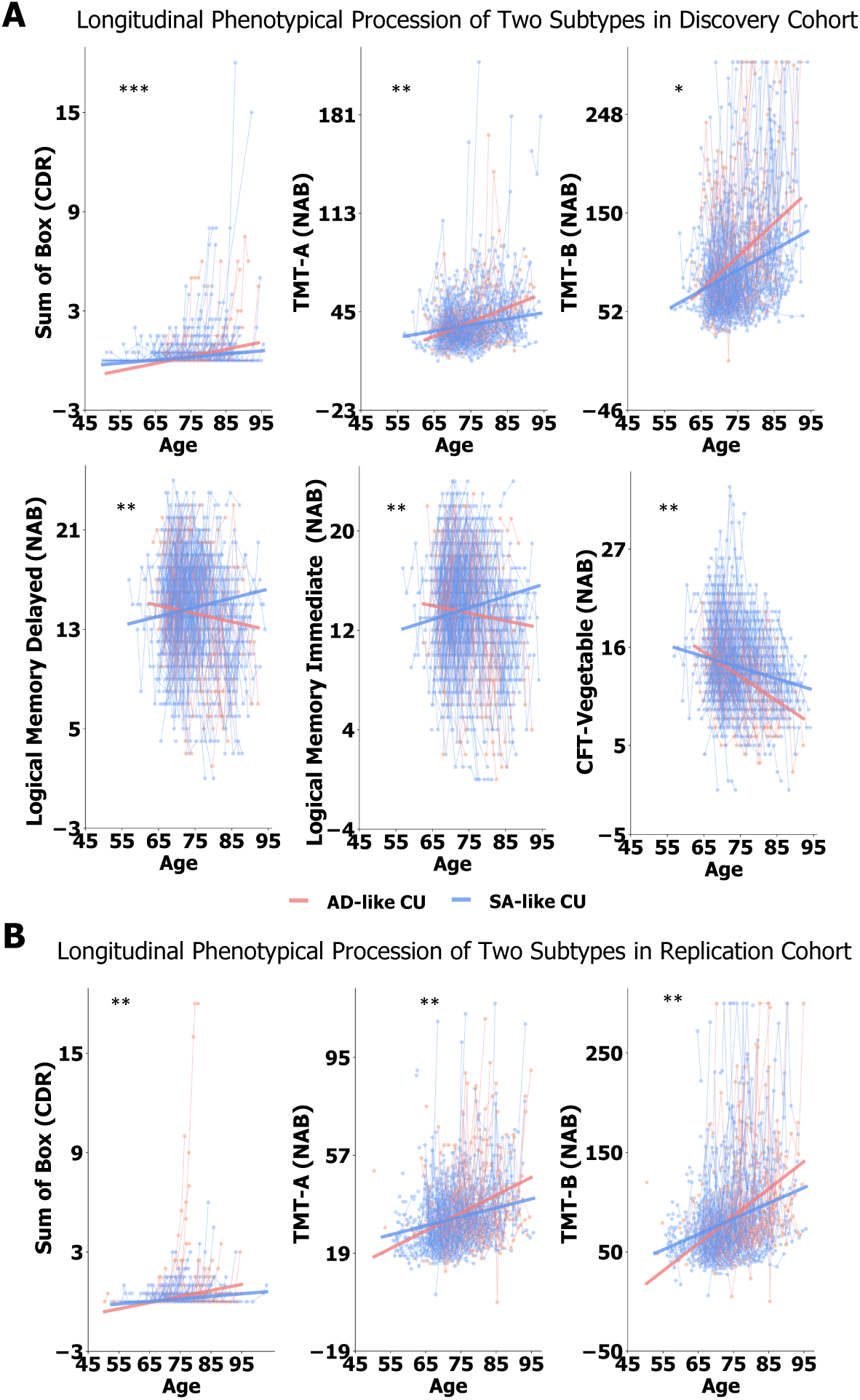
Longitudinal trajectories of characteristic variables for SA-like and AD-like subtypes in both the discovery and replication cohorts. Linear mixed effect models were used to examine the age-by-subtype interaction effect on various phenotypic variables (excluding demographics). Significant age-by-subtype interaction effects are indicated, showing that this interaction significantly explains variance in the phenotypic trajectories. FDR correction was applied to all p-values. The significance was annotated by * (p_fdr_ ≤ 0.05), ** (p_fdr_ ≤ 0.01), *** (p_fdr_ ≤ 0.001). **(A)** Longitudinal trajectories in the discovery cohort. **(B)** Longitudinal trajectories in the replication cohort. Higher scores of CDR and TMT, and lower scores of WAIS Logical Memory and CFT, reflect greater cognitive decline.

To assess the validity and added value of ARC-based subtyping, we conducted a control analysis using conventional hierarchical clustering to identify subtypes from phenotypic variables. Cluster stability was evaluated using consensus clustering (Figure S10). Both silhouette scores and changes in the area under the cumulative distribution function curve suggested that two clusters represented the most stable solution. Although this phenotypic clustering approach yielded subtypes with distinct baseline profiles, it revealed substantially fewer FC differences compared to ARC-based subtypes. Critically, although these subtypes differed in baseline phenotypic characteristics (Table S6), they did not exhibit significant differences in longitudinal cognitive trajectories, estimated by linear mixed-effects models. Attempts to replicate the phenotypic clustering results in the replication cohort using Euclidean distance to the discovery cohort cluster centers failed to reproduce FC or longitudinal outcome differences. These findings suggest that subtyping based solely on characteristic variables is less robust, less generalizable, and less sensitive to connectome-level and longitudinal heterogeneity than the ARC-based approach.

### Neuropathological accumulation differentiating ARC-defined CU subtypes

Aβ plaques and neurofibrillary tangles are critical pathology markers of AD progression^43^, and their accumulation has been observed in some CU elderly. To investigate how these biomarkers differ across ARC-defined subtypes, we first compared baseline levels of standardized uptake value ratio (SUVR) — a widely used measure of biomarker expression extracted from positron emission tomography (PET) data — between the SA-like and AD-like subtypes using the Kruskal-Wallis test. No significant differences in Aβ or tau uptake levels were observed at baseline (Figure S11), indicating that, at this stage, both subtypes exhibited comparable levels of amyloid and tau deposition. To explore potential differences in biomarker trajectories over time, we applied linear mixed-effects models to examine the age-by-subtype interaction effects on global Aβ and tau SUVRs. While no significant interaction was found for tau accumulation, Aβ demonstrated a distinct accumulation pattern, progressing more rapidly in the AD-like subtype compared to the SA-like subtype (Figure 5). To further localize this divergence, we analyzed regional Aβ deposition patterns across the two subtypes. As shown in Figure S12, the AD-like subtype displayed pronounced Aβ accumulation, particularly in the frontal and temporal lobes.

**Figure 5.**
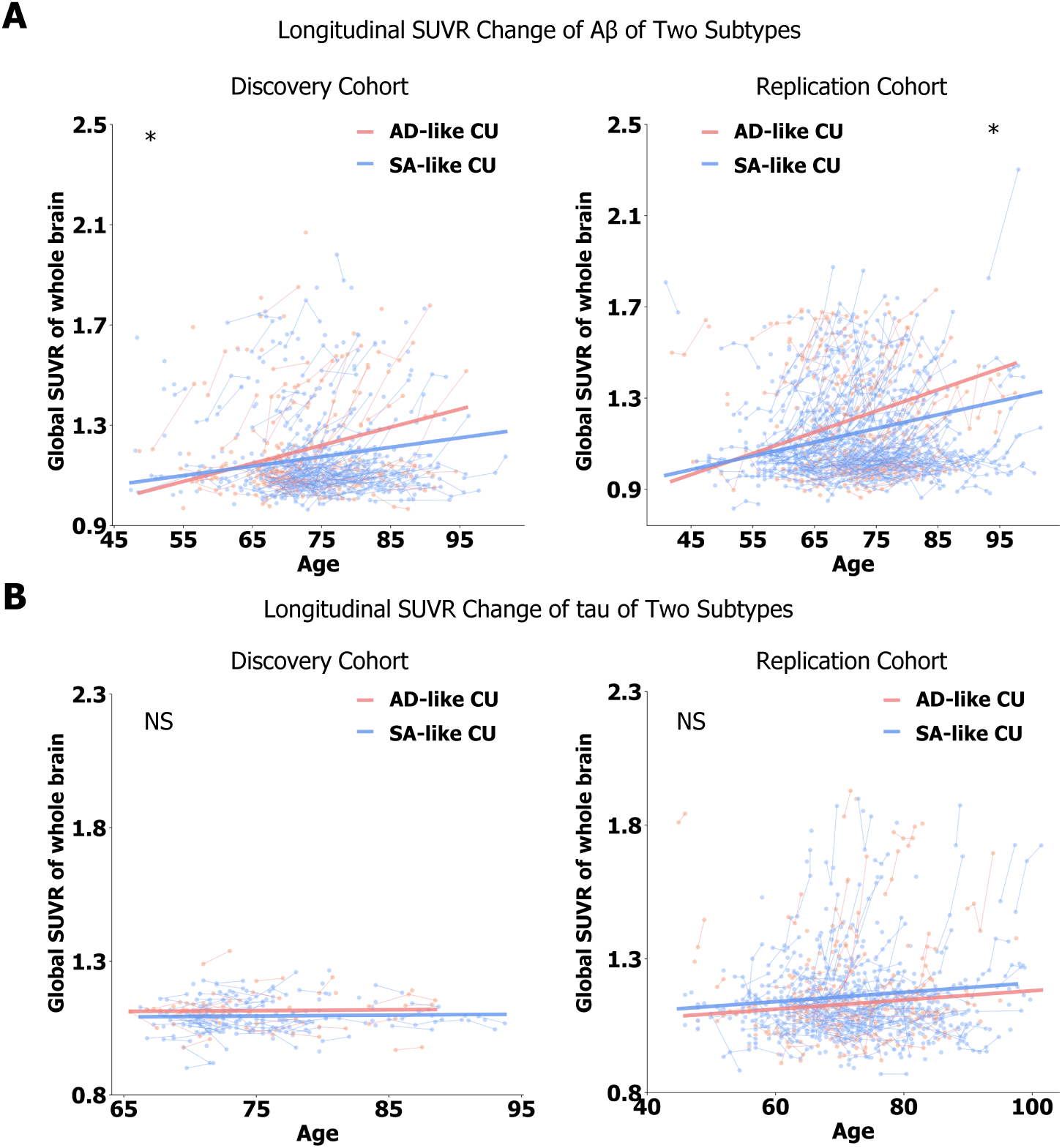
Longitudinal trajectories of globally expressed neuropathological biomarkers across the whole brain in SA-like and AD-like subtypes in the discovery and replication cohorts. Standardized uptake value ratio (SUVR) which accounts for variability in radiotracer delivery and clearance, was used to assess Aβ and tau pathology. A linear mixed-effects model was applied to examine the age-by-subtype interaction effect on biomarker expression. The significance was annotated by * (p ≤ 0.05), NS (No significance). **(A)** Longitudinal procession of Aβ between two subtypes. **(B)** Longitudinal procession of tau between two subtypes.

### Brain aging differentiates CU subtypes

The next key question we aimed to answer was whether baseline neurophysiological features could account for the distinct FC patterns and divergent longitudinal phenotypes observed between SA-like and AD-like CU subtypes. Among all examined characteristic variables, only chronological age significantly differed between the two subtypes at baseline (Tables S3-S5). This raised a key question: How can a difference in chronological age alone account for the marked divergence in long-term cognitive decline and Aβ accumulation between these CU subtypes? Chronological age is typically correlated with brain age, a metric that reflects neurobiological aging more directly^36,44^. Given this observation, we hypothesized that brain age, instead of chronological age, as a more precise indicator of neurobiological aging, would better differentiate the subtypes and more closely reflect their distinct cognitive trajectories. To test this, we applied our established brain age prediction model to CU individuals to estimate their brain age. For each CU individual, the ARC score was also calculated by z-scoring the weighted sum of FC features that differentiated the two subtypes based on the SA-AD classifier weights. We then examined the correlations between ARC scores, brain age, and chronological age. As shown in Figure S13, both brain age and chronological age were significantly correlated with ARC scores. However, evaluated by 1,000 bootstrapping trials, brain age exhibited a significantly stronger correlation with ARC scores than chronological age (p < 0.05 for both cohorts). Further analysis of BAG revealed a significant positive correlation with subtyping scores. Notably, AD-like CU individuals had significantly higher BAG than their SA-like counterparts, suggesting an accelerated brain aging process in the AD-like subtype.

### Genotypic profiles differentiating CU subtypes

To investigate the genetic basis underlying the phenotypic differences between the SA-like and AD-like CU subtypes, we conducted a GWAS using linear regression. As shown in Figure 6, three genetic loci were identified as significantly associated with subtype membership: rs2258615 (odds ratio [OR] = 0.38, p = 3×10^−6^), rs4772249 (OR = 0.44, p = 3×10^−6^), and rs17586545 (OR = 0.21, p = 5×10^−6^). These significant single nucleotide polymorphisms (SNPs) are primarily located in the genes *CLYBL* and *FRMD6* (Figure S14). The genetic variations in these loci may play a role in shaping the differential neurobiological and cognitive profiles observed between the subtypes.

**Figure 6.**
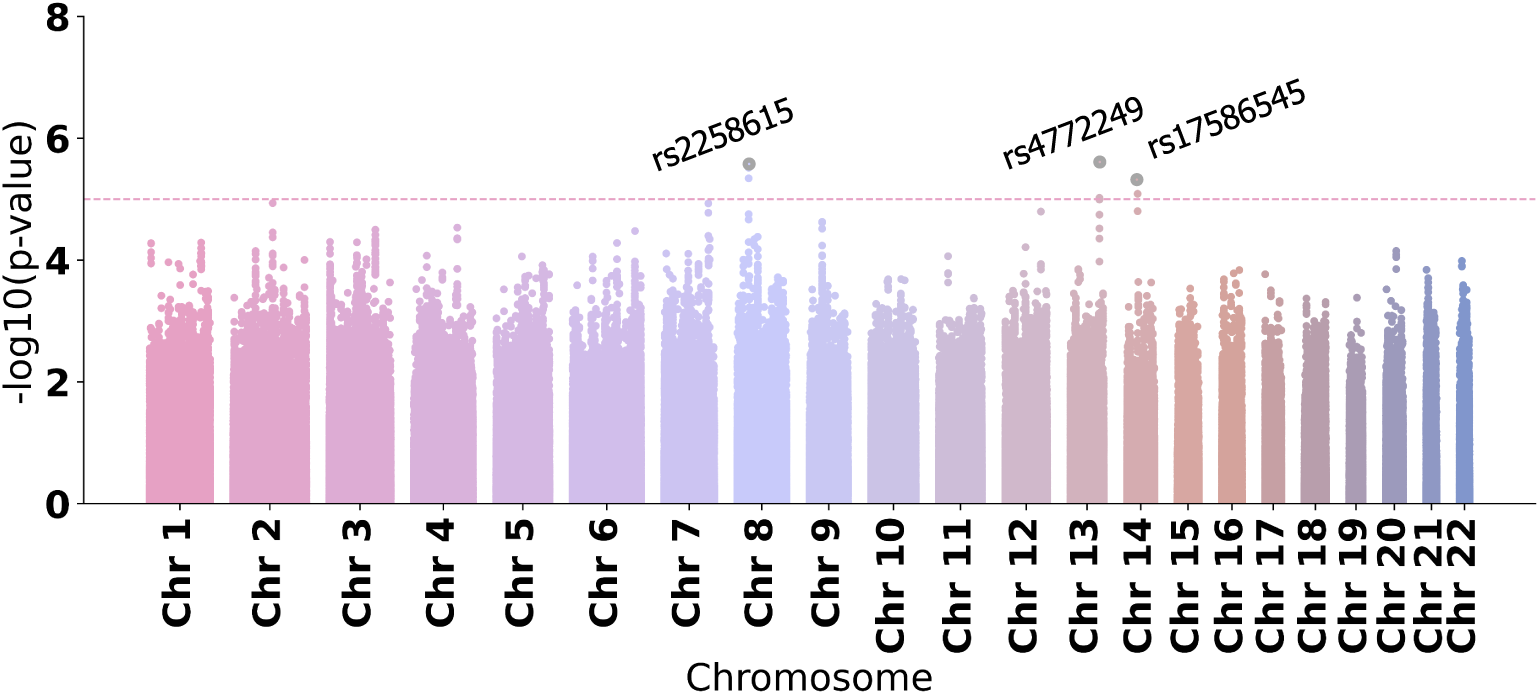
Manhattan plot of GWAS result associated with the differentiated FCs between AD-like (n = 151) and SA-like (n = 355) subtypes, using a logistic regression model with age, sex, and top five principal components^45,46^ accounting for population stratification as covariates. The red horizontal dashed line indicates the genome-wide significant threshold (p = 1 × 10^−5^).

## Discussion

Our study establishes FC as a sensitive and generalizable modality for identifying neurobiological signatures of cognitive resilience and vulnerability in aging. Leveraging machine learning, we developed a robust classifier that reliably distinguished SA from individuals with AD, prior to significant amyloid accumulation, with strong reproducibility in an independent replication cohort. The derived FC signature (ARC) was centered on the bilateral prefrontal cortex, angular gyrus, insula, and temporal poles, regions crucial for memory and cognitive control. When applied to CU individuals, the ARC signature stratified them into SA-like and AD-like subtypes. Despite comparable baseline cognitive performance, these subtypes exhibited distinct FC patterns, brain aging profiles, and longitudinal trajectories. Specifically, the SA-like subtype exhibited preserved cognitive function, lower risk of progression to MCI or AD, and reduced Aβ increases, particularly in the prefrontal and temporal cortices. Further, GWAS identified genetic variants in *CLYBL* and *FRMD6* that differentiate these subtypes. Collectively, these findings demonstrate that ARC detects biologically meaningful distinctions in brain aging before the onset of clinical symptoms, offering a powerful framework for early identification of cognitive resilience and targets for intervention.

The ARC signature delineated a network linking the prefrontal cortex, insula, and superior temporal regions, which collectively support executive functions, memory, and emotional regulation. Specifically, the prefrontal cortex and insula are primarily involved in cognitive control and working memory^47–51^, while temporal regions play essential roles in episodic and semantic memory^52^. Lower FCs involved in these regions were associated with cognitive resilience in SA^53–55^, whereas higher FCs were linked to memory decline and executive dysfunction in AD^56,57^. This is in line with the positive correlation between ARC scores with memory impairment and executive deficits measured by CDR and TMT, in our study. Furthermore, the ARC scores positively correlated with neuropsychiatric symptoms such as depression and apathy, suggesting its involvement in emotional regulation, motivational and goal-directed behaviors. This convergence aligns with our previous work linking memory deficits to affective disturbances in aging populations^42^. Together, these results suggest that disruption within the ARC network contributes to the complex clinical presentation of AD across cognitive, affective and motivational domains.

Beyond identifying ARC signature distinguishing SA and AD, our findings delineate two elder subgroups with distinct brain aging trajectories—resilient and vulnerable—reflecting resilience or vulnerability to neurodegeneration. The resilient trajectory, characterized by baseline SA-like FC patterns, was associated with preserved functional network integrity, long-term cognitive stability, and minimal neuropathological accumulation, suggesting inherent protective mechanisms against cognitive decline. In contrast, the vulnerable trajectory exhibits early FC disruption prior to clinical symptoms, serving as a preclinical marker of impending pathology, followed by faster Aβ deposition and cognitive deterioration, ultimately increasing MCI/AD risk. These findings position ARC as a sensitive biomarker for stratifying aging trajectories, enabling early identification of at-risk individuals and opportunities for targeted interventions to enhance cognitive resilience or slow pathological progression.

Interestingly, SA-like individuals with resilient aging trajectory showed better performance on the Logical Memory test with increasing age. While this may appear counterintuitive, it does not conflict with the broader pattern of decline observed in global CDR scores and TMT performance. Episodic memory, especially when tested under structured, low-distraction conditions, can remain stable, or even show modest improvements, may be attributed to practice effects or the influence of cognitive reserve, both of which are well-documented in CU older adults and individuals with early-stage MCI ^58–60^. In contrast, functional decline in daily activity like social communication—captured via clinician-rated and informant-based measures such as the CDR—can emerge earlier and may precede episodic memory impairments on neuropsychological assessments^61,62^. This dissociation underscores the limitations of relying solely on domain-specific cognitive tests and highlights the importance of multimodal assessments that integrate objective and informant-reported data to better capture subtle and heterogeneous aging trajectories^63^.

Notably, ARC-associated FC alterations were selectively linked to Aβ burden, but not tau, emphasizing key differences in the temporal dynamics of AD neuropathology. Aβ accumulation typically begins early and spreads diffusely across cortical association networks, while tau pathology emerges later and progresses along a more anatomically restricted trajectory^43,64,65^. This dissociation likely accounts for the closer correspondence between large-scale FC disruptions and Aβ accumulation during early disease stages. Notably, the spatial distribution of ARC pattern—characterized by increased connectivity in the superior prefrontal cortex and decreased connectivity in the temporal lobe—closely overlapped with regions exhibiting amyloid pathology. The anatomical co-localization of ARC-defined FC abnormalities and Aβ burden suggests a mechanistic axis linking network dysfunction to molecular pathology.

Importantly, previous studies have shown that cortical organization across anatomical, functional, evolutionary, and molecular domains—including gene expression and neuronal differentiation— converges along a shared brain developmental axis^66^. Analogously, inspired by this convergence, our findings, together with earlier reports in elderly population^67–70^, that observed co-occurrence of FC disruption and Aβ deposition in the same cortical areas, support the existence of a brain aging axis. This proposed axis reflects a multimodal vulnerability gradient, where spatially aligned functional and pathological disturbances interact to drive individual trajectories of neurodegenerative risk. Future research integrating brain patterns of gene expression and neuron-type-specific differentiation may further verify this brain aging axis.

Our GWAS identified two novel loci, *CLYBL* and *FRMD6*, differentiating individuals along the vulnerable versus resilient aging trajectories. Although these genes have been relatively understudied in neurodegeneration, emerging evidence suggests their involvement in processes critical to aging. *CLYBL* regulates citrate metabolism and inflammatory pathways^71^, while *FRMD6* modulates neuronal morphogenesis and synaptic architecture^72,73^—both implicated in neurodegenerative vulnerability^74,75^. These findings suggest that neuroinflammation and synaptic dysfunction may contribute to individual differences in brain aging resilience. Further work is needed to elucidate the mechanistic roles of these loci and to explore their potential as therapeutic targets for promoting healthy brain aging.

Several limitations warrant consideration. First, although our sample was relatively large and multi-site, expanding analyses to more diverse ethnic groups will be essential for enhancing the generalizability of our findings. Second, the use of resting-state FC alone may overlook other critical neurobiological features. Integrating multi-modal neuroimaging measures, such as structural connectivity and structure-function coupling, may offer additional understanding of aging-related brain changes.

Finally, while our GWAS identified promising genetic associations, larger replication studies will be essential to validate the roles of these loci in modulating brain aging trajectories and resilience. In summary, our study established FC as a sensitive and generalizable marker of cognitive resilience and vulnerability across the aging spectrum. Using a data-driven machine learning framework, we identified a robust ARC signature involving the prefrontal, temporal, and insular cortices that reliably distinguished SA from individuals with AD across replication cohorts. This FC signature captured a reproducible neurofunctional resilience phenotype, characterized by preserved network architecture and youthful brain aging trajectories despite chronological aging. When applied to CU individuals, the ARC signature stratified participants into biologically distinct subtypes. Although baseline cognition was comparable between groups, longitudinal analyses revealed that the AD-like subtype exhibited faster cognitive decline, greater amyloid-β accumulation, and increased risk of clinical progression, whereas the SA-like subtype demonstrated cognitive stability and delayed pathological burden. These findings position the ARC signature as an early indicator of vulnerable versus resilient brain aging, preceding both clinical symptoms and neuropathological changes. Moreover, GWAS implicated *CLYBL* and *FRMD6* as genetic modulators of these trajectories, highlighting potential links between neuroinflammatory and neuronal morphogenesis pathways and aging resilience. Together, our results advance a mechanistic model in which the preservation of functional network organization confers protection against neurodegeneration, offering a novel framework for early risk stratification and paving the way for targeted interventions aimed at sustaining cognitive health aging population.

## Methods

### Discovery cohort

#### OASIS-3 dataset

In this study, we utilized the longitudinal data collected from the Open Access Series of Imaging Studies (OASIS-3) database, which enrolled participants at Washington University in St. Louis over 15 years ^76^. 1098 participants aged between 42 and 95 were enrolled beginning in 2005. All participants in OASIS-3 provided informed consent in accordance with procedures approved by the Institutional Review Board of Washington University School of Medicine. The dataset comprised resting-state fMRI scans and clinical assessments as well as neuropsychological measures. Exclusion criteria included medical conditions that precluded longitudinal participation (e.g. end-stage renal disease requiring dialysis) or medical contraindications for the study arms (e.g. pacemaker for MRI, anticoagulant use for lumbar puncture).

#### HABS dataset

The Harvard Aging Brain Study (HABS) was launched in 2010^77^. 290 cognitively healthy individuals older than 65 were registered. As a longitudinal project, HABS cohort collected the neuropsychological measurements and neuroimages of participants each year until 2015.

### Replication cohort and genotypic dataset

#### ADNI dataset

The Alzheimer’s Disease Neuroimaging Initiative (ADNI) is a longitudinal observational program, which enrolled 2433 participants aged 55 to 90 years, aiming to collect and analyze biomarkers related to the progression to AD^78^. Launched in 2003, ADNI has collected serial MRI and neuropsychological measures. The participant recruitment for ADNI is approved by the Institutional Review Board of each participating site. Exclusion criteria comprised current use of psychoactive medication, a history of schizophrenia, substance use and various mental disorders^78^. Additionally, ADNI dataset was used as a genotypic dataset, since SNP was only available in this cohort.

#### Neuropsychological assessments

Demographic variables were analyzed, including age, sex, years of education, and race. To assess dementia symptoms, *Clinical Dementia Rating Scale* (CDR) was used^79^. Specifically, it evaluates cognitive and functional performance in six domains: memory, orientation, judgment and problem solving, community affairs, home and hobbies, and personal care. Verbal memory, learning and recall were assessed with the *Rey Auditory Verbal Learning Test* (RAVLT)^80^, in which participants recalled a list of 15 unrelated words over five learning trials, followed by an interference list and a delay before final recall. Derived RAVLT scores offered insights into distinct cognitive processes^81,82^, including RAVLT Immediate highlighting total learning ability, RAVLT Learning emphasizing learning rate, and RAVLT Percent Forgetting emphasizing forgetting rate or delayed memory. The functional ability is evaluated by *Functional Activities Questionnaire* (FAQ) ^83^ in 10 daily activities: 1) writing checks, paying bills, and keeping financial records; 2) assembling tax records and making out business and insurance papers; 3) shopping alone for clothes, household necessities, and groceries; 4) playing a game of skill such as bridge, other card game, or chess; 5) heating water for coffee or tea and turning off the stove; 6) preparing a balanced meal; 7) keeping track of current events; 8) paying attention to and understanding a TV program, book, or magazine; 9) remembering appointments, family occasions, and medications; and 10) travel out of the neighborhood. Moreover, the *neuropsychological assessment battery (NAB)* ^34^ was applied in assessing neuropsychiatric disturbances and cognitive and behavioral dysfunction. For the NAB, ten neuropsychological tests measure attention/working and episodic memory, executive function, and language, including the *Digit Span Forward* (DIGIF), *Backward test* (DIGIB), and *Logical Memory in Wechsler Memory Scale* (WAIS-R)^84^, *Category fluency test* of animal and vegetable (CFT-ANI/ CFT-VEG)^85^, *Trail Making Test* Part A and B^86^, and *Boston Naming Test*^35^.

#### Superager definition

The definition of SA in this study was based on cognitive resilience in aging, integrating episodic memory and executive function criteria. The definition encompassed three key aspects: an advanced age threshold (≥ 75 years), superior episodic memory performance compared to middle-aged adults (under 65), and executive function comparable to age-matched older adults^10,25^. Specifically, SA was required to exhibit episodic memory performance exceeding the mean of middle-aged adults and executive function at least one standard deviation above the mean of age-matched normal elderly individuals. In the primary analysis, executive function was assessed using the TMT-B, with a cutoff score of ≤166^29^ applied consistently across cohorts. Episodic memory evaluation utilized the RAVLT delayed recall (≥ 9) in the replication cohort, a well-validated measure in SA research^25,87^. In the discovery cohort where RAVLT was unavailable, WAIS-R Logical Memory (IA ≥13 and IIA ≥11)^26,27^ served as an alternative based on its established correlation with RAVLT performance^88,89^ and prior use in SA studies^18,90,91^. To validate the robustness of our further analysis based on classification between SA and AD, we implemented two additional alternative strategies in defining SA. In one approach, SA criteria in the discovery cohort remained unchanged, but in the replication cohort, WAIS-R Logical Memory replaced RAVLT to maintain consistency in memory measures across cohorts (Figure S15). The second approach expanded the executive function assessment by incorporating the ADNI-EF composite measure^92^, which integrates multiple cognitive domains including animal and vegetable naming (CFT-ANI/VEG), processing speed (TMT-A/B), and working memory (WAIS-R Digit Symbol) following published SA studies^11,25,93–95^. This alternative approach defined executive function in the ADNI dataset as ADNI-EF ≥ −0.02. In the OASIS-3 and HABS datasets, executive function was assessed using CFT-ANI (≥ 14), CFT-VEG (≥ 15)^96^, TMT-A (≤ 68), TMT-B (≤ 166)^92^, and WAIS-R Digit Symbol (≥ 24.5)^97^. The framework involving such an approach in defining SA was shown in Figure S16. The results of classification between AD and SA defined using various approaches were mentioned in “*Classification between SA and AD*” section of method.

#### MRI acquisition and preprocessing

MRI data in OASIS-3 were scanned in three different Siemens scanners (Siemens Vision 1.5 T, 2 scanners of TIM Trio 3T) and Siemens BioGraph mMR PET-MR 3T. High resolution T1-weighted structural image (TR = 2.4 s, TE = 3.08 ms, FOV = 256 × 256 mm, FA = 8°, voxel size 1 × 1 × 1 mm^3^) and resting-state functional image (TR = 2.2 s, TE = 27 ms, FOV = 240 × 240 mm, FA = 90°, duration = 6 min, voxel size 4 × 4 × 4 mm, 36 slices) were obtained. MRI data in HABS were collected at the Massachusetts General Hospital Martinos Center for Biomedical Imaging in Charlestown, using 3T Siemens Tim Trio 3T scanners with a 12-channel phased-array head coil^77^. Some parameters of fMRI scans are: TR = 2 s, FA = 85°, slice thickness = 3 mm. MRI data in ADNI were scanned in 3T Philips system using magnetization-prepared rapid acquisition gradient echo^78^. Some scanner parameters of resting-state fMRI are: TR = 6 s, TE = 32 ms, FA = 50°, slice thickness = 2.5 mm.

The acquired resting-state fMRI data were preprocessed using the reproducible fMRIPrep pipeline ^98^. Several important steps are summarized as follows: (1) The T1 weighted image underwent correction for intensity nonuniformity and skull stripping. (2) Spatial normalization was achieved through nonlinear registration, utilizing the T1w reference ^99^. (3) FSL was employed to segment brain tissue, including cerebrospinal fluid, white matter, and grey matter, from the reference, brain-extracted T1 weighted image ^100^. (4) Fieldmap information corrected distortion in low- and high-frequency components due to field inhomogeneity. (5) The blood-oxygen-level-dependent (BOLD) reference was transformed to the T1-weighted image using a boundary-based registration method with nine degrees of freedom to address remaining distortion ^101^.

#### Calculation of resting-state FC

The BOLD signals of preprocessed fMRI were averaged into time series of 100 regions of interest (ROIs) defined by the Schaefer parcellation ^30^. Pearson correlation was then computed between time series of each pair of ROIs, resulting in 4950 FCs for each participant. Fisher’s r-to-z transformation was applied to enhance normality of connectivity, followed by z-score normalization.

#### PET acquisition, preprocessing and postprocessing

The standardized uptake value ratio (SUVR) data extracted from preprocessed Pittsburgh Compound B (^11^C-PIB) PET images in OASIS-3, HABS and ADNI datasets were downloaded from their official websites of ^102–104^. ^11^C-PIB PET images were widely used to explore the expression of amyloid-β deposits in the brain^105^. The details of acquisition parameters, preprocessing and postprocessing steps can be found on the websites^106^ and in the eMethod section in our supplement. SUVR of each ROI defined in the Desikan atlas^107^ was then calculated. The global SUVR were averaged from the SUVR of all ROIs.

#### Genotype processing and quality control

Genotypic data was only available in the ADNI cohort. DNA was extracted from blood on genotyping array of three illumine platform: Human610-Quad, HumanOmniExpress, and Omni 2.5M. Quality control was performed in several standard procedures^22,108,109^, including removal of single nucleotide polymorphisms (SNPs) and samples with > 5% genotype missingness, removal of SNPs with < 1% minor allele frequency (MAF) or Hardy-Weinberg Equilibrium (HWE) p-values < 10^−6^, and removal of samples with sex discrepancies, cryptic relatedness (pi-hat > 0.25). Then genotypes were imputed using Minimac4 locally based on a precompiled 1000 Genomes reference panel^110^. Post-imputation quality control steps included removal of SNPs with imputation quality score R^2^ < 0.90, call rate < 95%, MAF < 1%, or HWE p-value < 10^−6^. Importantly, two previously identified APOE SNPs important in AD susceptibility (rs429358, rs7412) were not available on the Illumina array. Following the reference^111^, we determined the genotypes of the two APOE SNPs (rs429358, rs7412) using the APOE ε2/ε3/ε4 status information from the ADNI clinical records for each participant. A total of 3417996 SNPs remained.

#### Classification between SA and AD

A linear support vector machine (SVM) was employed to classify SA and AD patients. To reduce feature dimensionality, we first applied two-sample t-tests to identify FC that significantly differed between groups (p < 0.05). FC selection was performed exclusively on the training set, and the resulting feature mask was applied to the test set for classification. L1 regularization was used in SVM to further avoid overfitting with a penalty parameter 0.1. To enhance model robustness, we increased the training sample size by incorporating multiple runs and sessions per participant and weighted more for the group with smaller sample size. We evaluated classification performance through 10-fold cross-validation. The ARC was characterized by the t-values of FCs that significantly differentiated SA and AD and contributed to classification with nonzero weights. To further validate the generalizability of the identified FC signature, ten SVM classifiers from cross-validation runs were applied to an independent replication cohort. Predicted labels were determined by averaging classification probabilities across models, followed by performance evaluation using metrics including accuracy, sensitivity, specificity, and AUC. Since SA was defined by an age threshold of ≥75 years, we selected age-matched AD participants for classification. Demographic details for the discovery cohort (SA: 107 female [59%]; median [IQR] age, 80 [77-82] years; AD: 89 female [80%]; median [IQR] age, 80[77-83] years) and replication cohort (SA: 42 female [64%]; median [IQR] age, 79 [77-83] years; AD: 32 female [41%]; median [IQR] age, 80 [77-85] years) are summarized in Tables S1–S2.

To validate measurement equivalence between memory assessments, we applied classifiers trained on the discovery cohort to the replication cohort with SA defined by WAIS-R instead of RAVLT (Figure S15). This approach yielded comparable performance (Figure S17A: accuracy=0.70, sensitivity=0.70, specificity=0.71, AUC=0.74) to our primary analysis using RAVLT-defined SA (Figure 1A). The substantial overlap (> 50%) between SA identified by both definitions likely explains this consistency in classification performance (Figure S17B). Furthermore, prior studies have defined SA using a more comprehensive assessment of executive function, incorporating CFT-ANI, CFT-VEG, TMT-A, TMT-B, and WAIS-R Digit Symbol^11,25,93–95^. To align with these approaches, we adopted this broader criterion for defining SA (Figure S16) and repeated the classification procedures outlined in Figure S2. The resulting classification performance and FC signatures, as shown in Figure S18, remained consistent with those obtained in our main analysis (Figure 1). These findings reinforce the robustness of the predictive FC signature across different SA definitions.

#### Subtyping analyses on CU individuals

We applied all cross-validated models to CU individuals at baseline to stratify them into SA-like and AD-like subgroups. Specifically, we determined SA-like CU subtype if the participants were predicted as SA in more than half of the cross-validation folds, while others were classified as AD-like CU. Then, we compared FCs between identified subtypes, using linear regression, with subtype label as the main factor and age and sex as covariates. FDR correction was used for significantly differentiated FCs. The same step was used to detect the difference of SUVR of whole brain between two subtypes. The 2-tailed χ^2^ test, Kruskal-Wallis analysis, were used to detect differences in phenotypical variables between two subtypes. We used FDR to correct the significance of comparison results of all phenotypical variables including demographic information, each item of CDR, NAB, and FAQ. To validate the longitudinal cognitive abnormal progression between identified subtypes, we employed linear mixed-effect models. These models included the phenotypical variable at each study visit as the dependent variable, while subtype label, age, and the interaction of subtype label and age as independent variables. We applied FDR correction to the p-values of interaction effects across all phenotypic items. Additionally, linear mixed-effect models were used to investigate the change of expression SUVR of Aβ and tau of whole brain and all regions over time, with subtype label, age, and the interaction of subtype label and age as independent variables. All these statistical analyses were reproduced in the replication dataset.

We finally conducted GWAS to explore the association between genotype and differentiated FCs of two subtypes, using linear regression in PLINK (version 2.0) to analyze the SNP information from ADNI cohort. In the regressor, imputed allele dosages were utilized as independent variables and included adjustments for sex, age and the top five principal components. Statistical significance was defined as *α* = 1 × 10^−5^, a widely used suggestive threshold in previous ADNI GWAS studies^22,108^.

## Acknowledgement

This work was supported in part by NIH grant nos. R21AG080425, R01MH129694, R01AG081768, and R21MH130956, Alzheimer’s Association Grant (AARG-22-972541), and Lehigh University FIG (FIGAWD35) and CORE grants. Portions of this research were conducted on Lehigh University’s Research Computing infrastructure partially supported by NSF Award 2019035. G.A.F. was also supported by NIH grant nos. R01MH132784, R01MH125886, philanthropic funding, and grants from the One Mind - Baszucki Brain Research Fund, the SEAL Future Foundation, and the Brain and Behavior Research Foundation.

## Author Contributions

K.Z. conceptualized and designed the work, wrote the code, analyzed and interpreted the data, and drafted and revised the manuscript. H.X., G.A.F., N.B.C., T.J., R.S.O., A.C., and F.V.L. refined the design of the work, interpreted the data, and revised the manuscript. Y.Z. conceptualized and designed the work, oversaw the analysis and interpretation of the data, and revised the manuscript.

## Data Availability

The OASIS-3 dataset is publicly available (https://www.oasis-brains.org/). The ADNI dataset is publicly available (https://adni.loni.usc.edu/). The HABS dataset is publicly available (https://habs.mgh.harvard.edu/researchers/data-details/).

## Supplement

### eMethod

#### PET preprocessing and postprocessing

In OASIS-3 cohort, participants underwent positron emission tomography (PET) on one of three different Siemens scanners: ECAT HRplus 962 PET scanner, Biograph 40 PET/CT scanner, Biograph mMR PET-MR^1^. Pittsburgh Compound B (^11^C-PIB) was used to investigate amyloid-β deposits in the brain. Participants received an I.V. bolus administration of 6 - 20 mCi of ^11^C-PIB with a 60-minute dynamic PET scan in 3D mode (24 × 5 sec frames; 9 × 20 sec frames; 10 × 1 min frames; 9 × 5 min frames). PET imaging was preprocessed using the PET Unified Pipeline (PUP) via XNAT^2^, which included steps for spatial smoothing (to 8mm resolution), inter-frame motion correction, and PET-to-MR registration using a vector-gradient algorithm. Regional PET processing was based on FreeSurfer segmentation, and voxel-wise standardized uptake value ratio (SUVR) images were generated^3^. The PUP pipeline accounted for partial volume effects using a regional spread function approach to improve quantification and sensitivity to longitudinal changes^4^. Two modeling approaches were applied: Logan graphical analysis for dynamic PET data to calculate binding potential (BPND) and non-Logan analysis for static data, with SUVR estimates calculated using specific tracer time windows.

In HABS cohort, participants underwent PET on a Siemens ECAT HR+ PET scanner (3D mode; 63 image planes; 15.2 cm axial field of view; 5.6 mm transaxial resolution; 2.4 mm slice interval). Before injection, 10-minute transmission scans for attenuation correction were collected. After injection of 8.5-15 mCi ^11^C-PIB, 60 minutes of dynamic data were acquired in 3D acquisition mode and reconstructed in 39 frames (8 × 15s, 4 × 60s, and 27 × 120s). For PET preprocessing, mean image (first 8 minutes post-injection for ^11^C-PIB) was generated and coregistered to the corresponding FreeSurfer-processed T1 image using 6 degrees rigid body registration with SPM12’s spm_coreg function. The gtmseg atlas, created with FreeSurfer version 6, was utilized with the mri_gtmpvc function to produce partial volume corrected images based on region-based voxel-wise correction method^5^. Regional SUVRs were calculated for the target ROIs and for the OFF regions from PVC images.

In ADNI cohort, participants underwent PET on a Siemens ECAT HR+ PET scanner (3D mode; 63 image planes; 15.2 cm axial field of view; 5.6 mm transaxial resolution; 2.4 mm slice interval). Before injection, 10-minute transmission scans for attenuation correction were collected. After injection of 8.5-15 mCi ^11^C-PIB, 60 minutes of dynamic data were acquired in 3D acquisition mode and reconstructed in 39 frames (8 × 15s, 4 × 60s, and 27 × 120s).

The PET preprocessing pipeline, relying on SPM12 and PETPVC, involves registering the PET image to its corresponding T1w image in native space using SPM’s Co-register method. An optional partial volume correction (PVC) step using the regional voxel-based method was applied with tissue maps from the T1w image. The PET image is then transformed into MNI space using the same T1w transformation (DARTEL to MNI), intensity normalized to a reference region (e.g., eroded cerebellumpons), and masked to exclude non-brain regions using tissue probability maps. The resulting SUVR images are in a common space, enabling voxel-wise correspondence across subjects.

**Figure S1.**
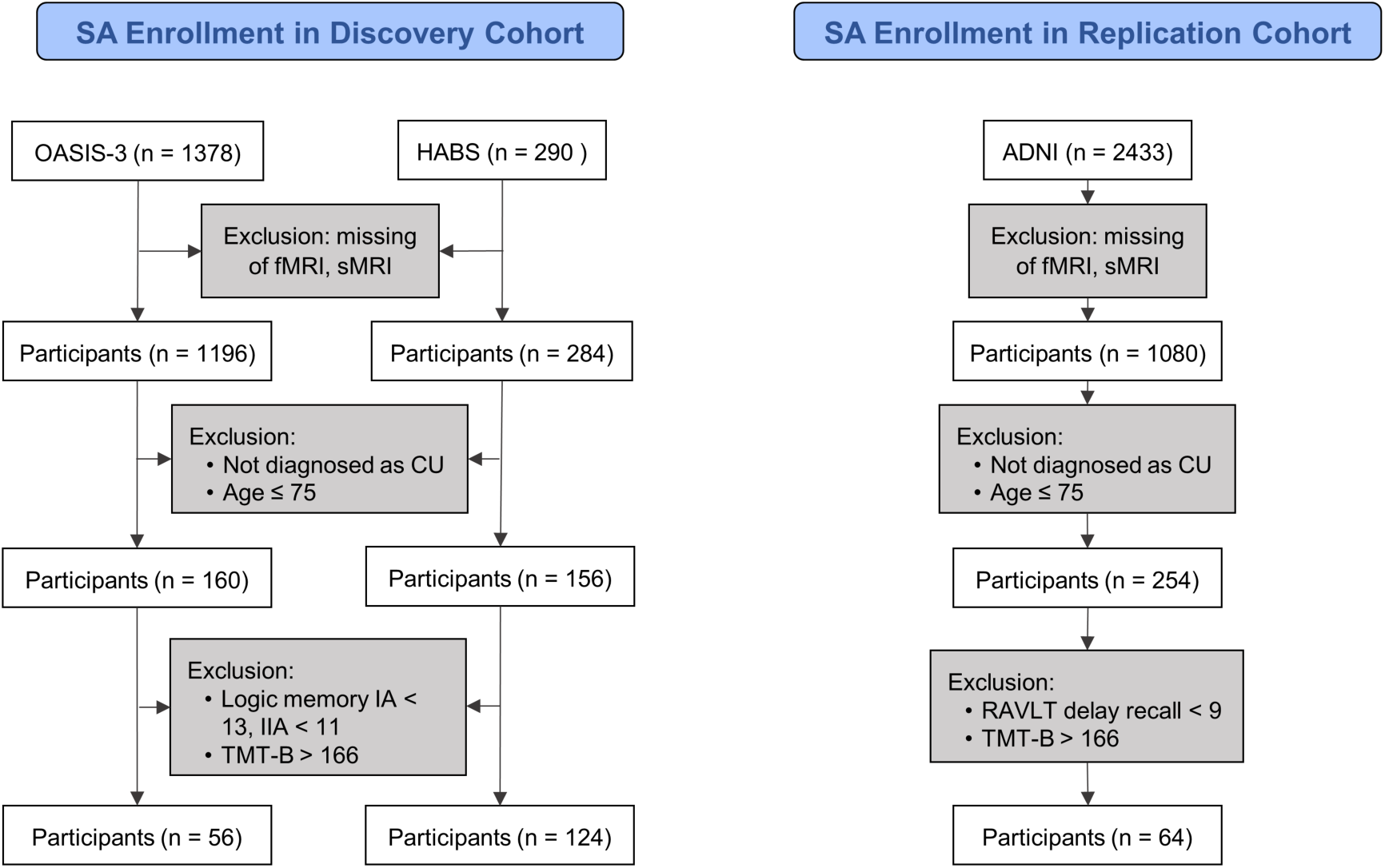
Flow diagram for SA inclusion in our study. SA were defined based on age and performance in episodic memory and executive function assessments. While most SA studies evaluate episodic memory using the Rey Auditory Verbal Learning Test (RAVLT) delayed recall score, the protocol of public dataset consisted of discovery cohort did not include RAVLT. As an alternative, we used the Wechsler Memory Scale Logical Memory test to assess episodic memory.

**Figure S2.**
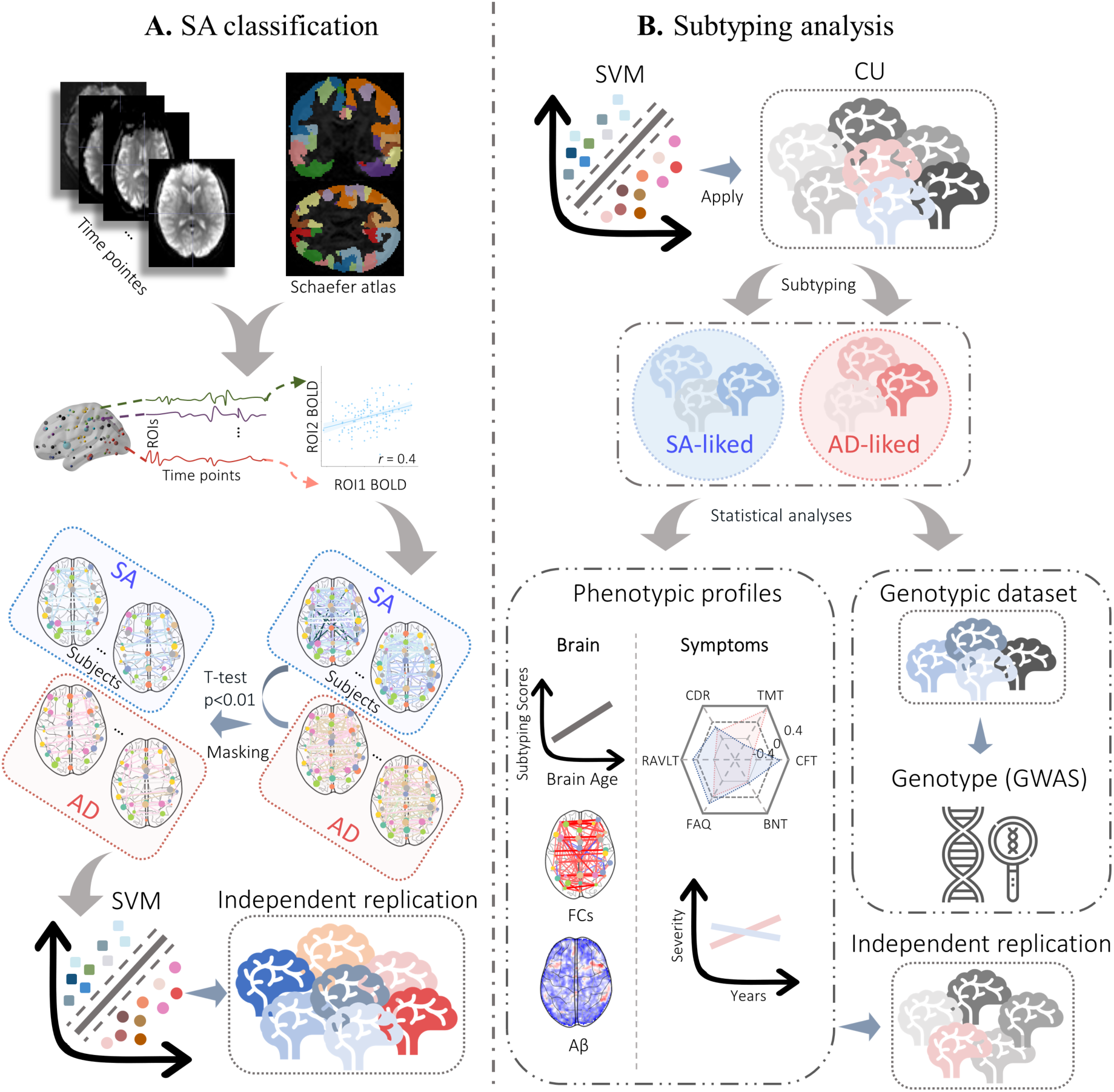
Study framework. **(A)** Classification between SA and AD. The Schaefer atlas was used to segment the brain into 100 regions, from which time series were extracted from functional magnetic resonance imaging data for all participants. FCs differentiated between SA and AD were identified using a two-sample t-test and subsequently used to train a support vector machine (SVM) classifier to distinguish these two groups. Model performance was assessed using ten-fold cross-validation within the discovery cohort and further validated in an independent replication cohort to confirm generalizability. **(B)** Subtyping analysis of cognitively unimpaired (CU) individuals. The well-trained classifiers were applied to group CU individuals into SA-like and AD-like subtypes based on their FCs. Comprehensive phenotypic profiles were obtained by comparing brain age FCs and cognitive performance at baseline between the subtypes. Longitudinal changes in amyloid-beta expression and cognitive decline were analyzed to assess subtype-specific trajectories. These findings were then reproduced in an independent replication cohort. Finally, a genome-wide association study (GWAS) was conducted on single nucleotide polymorphism (SNP) genotypes from the genotypic dataset to identify potential genetic contributors to the phenotypic differences between the subtypes.

**Figure S3.**
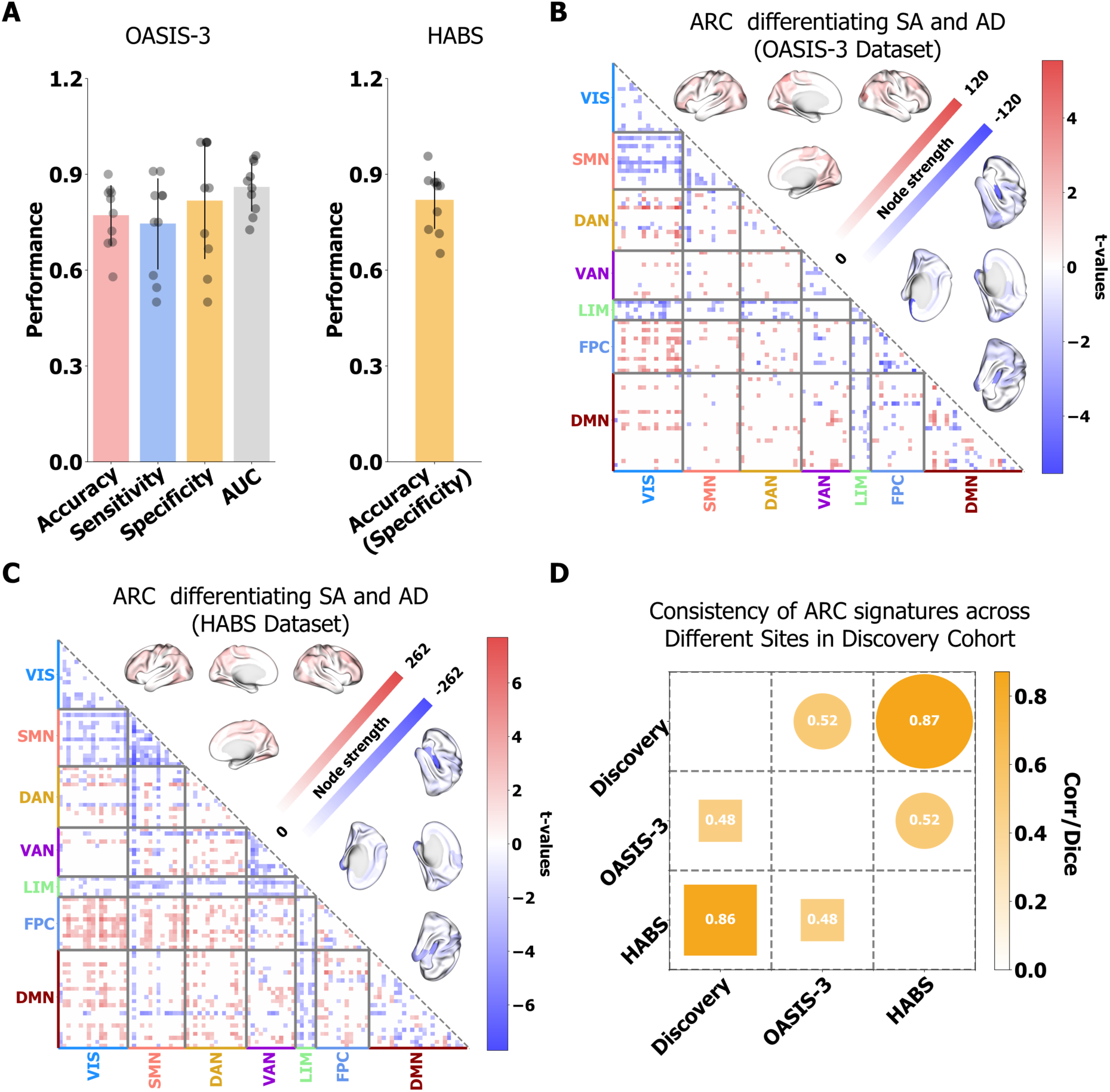
Cross-site classification in the discovery cohort. **(A)** Classification performance for each dataset consisted of the discovery cohort. For the HABS dataset, only accuracy (equal to specificity) is reported since these datasets do not contain participants with AD. **(B), (C)** ARC signatures that significantly differentiated SA from AD as detected by two-sample t-tests (FDR-corrected p < 0.05) in OASIS-3, and HABS datasets were evaluated, using two-sample t-tests (p < 0.05, false discovery rate (FDR) corrected) to compare FCs with non-zero weights in the classifiers distinguishing SA and AD. Hypo-connections (stronger in SA) are colored red, while hyper-connections (stronger in AD) are colored blue. Node strength, derived from summed positive and negative t-values of distinguishing FCs, is visualized on the brain surface. **(D)** Cross-site consistency of ARC signatures. Heatmaps display Kendall’s rank correlation and Dice coefficient. The upper triangular heatmap represents Kendall correlation coefficients (circle box), while the lower triangular heatmap represents Dice coefficients (rectangle box). All the values in the heatmap were significant (p < 0.001).

**Figure S4.**
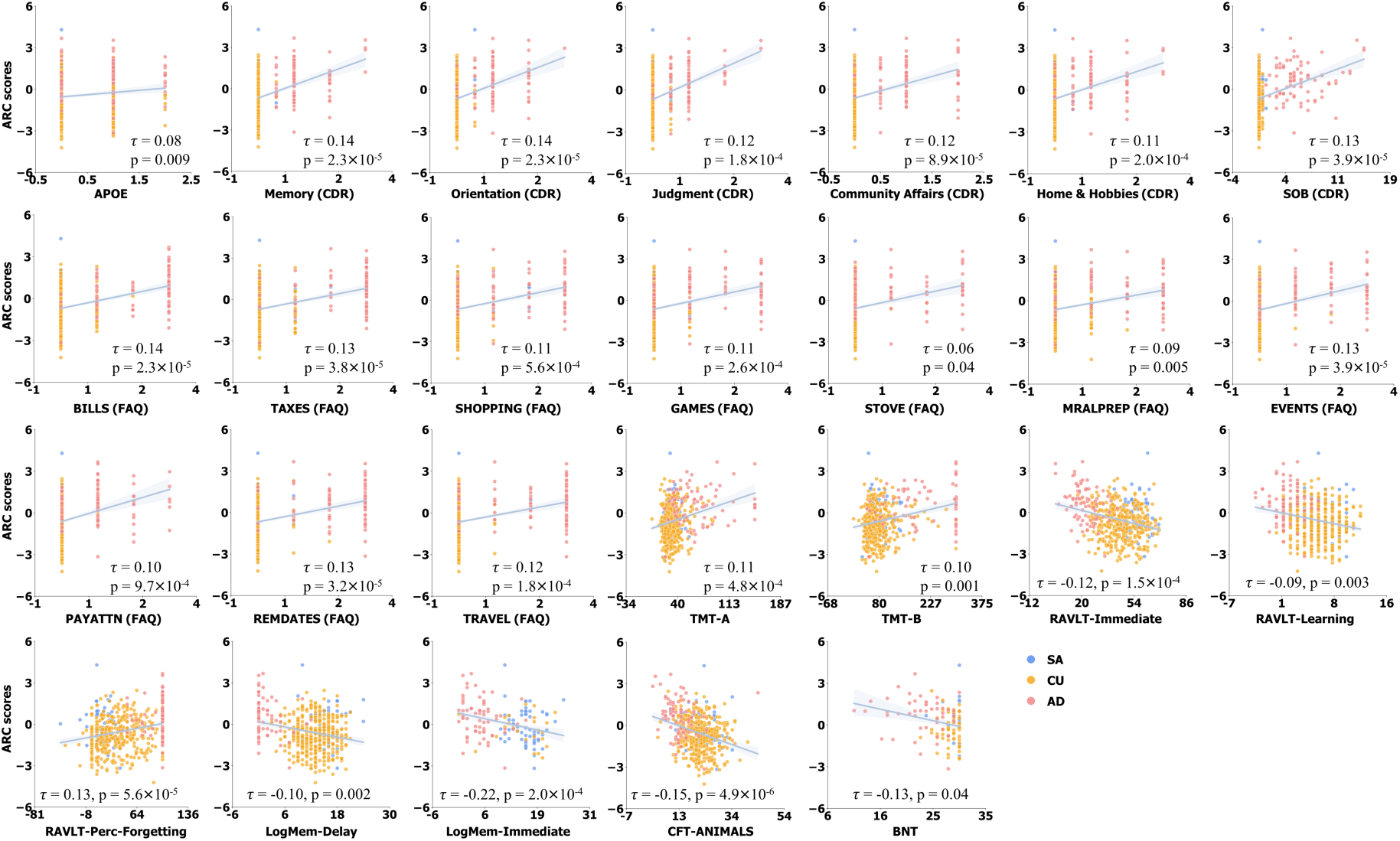
Phenotypic profiles correlated with ARC across SA, CU, AD. The correlation between various phenotypic variables (excluding demographic variables) and the ARC scores was examined across SA, CU, and AD in the replication cohort. Kendall rank correlation was used to assess relationships, controlling age since age of CU at baseline was younger than SA and AD. FDR correction was applied to all p-values. Only the significant results were shown.

**Figure S5.**
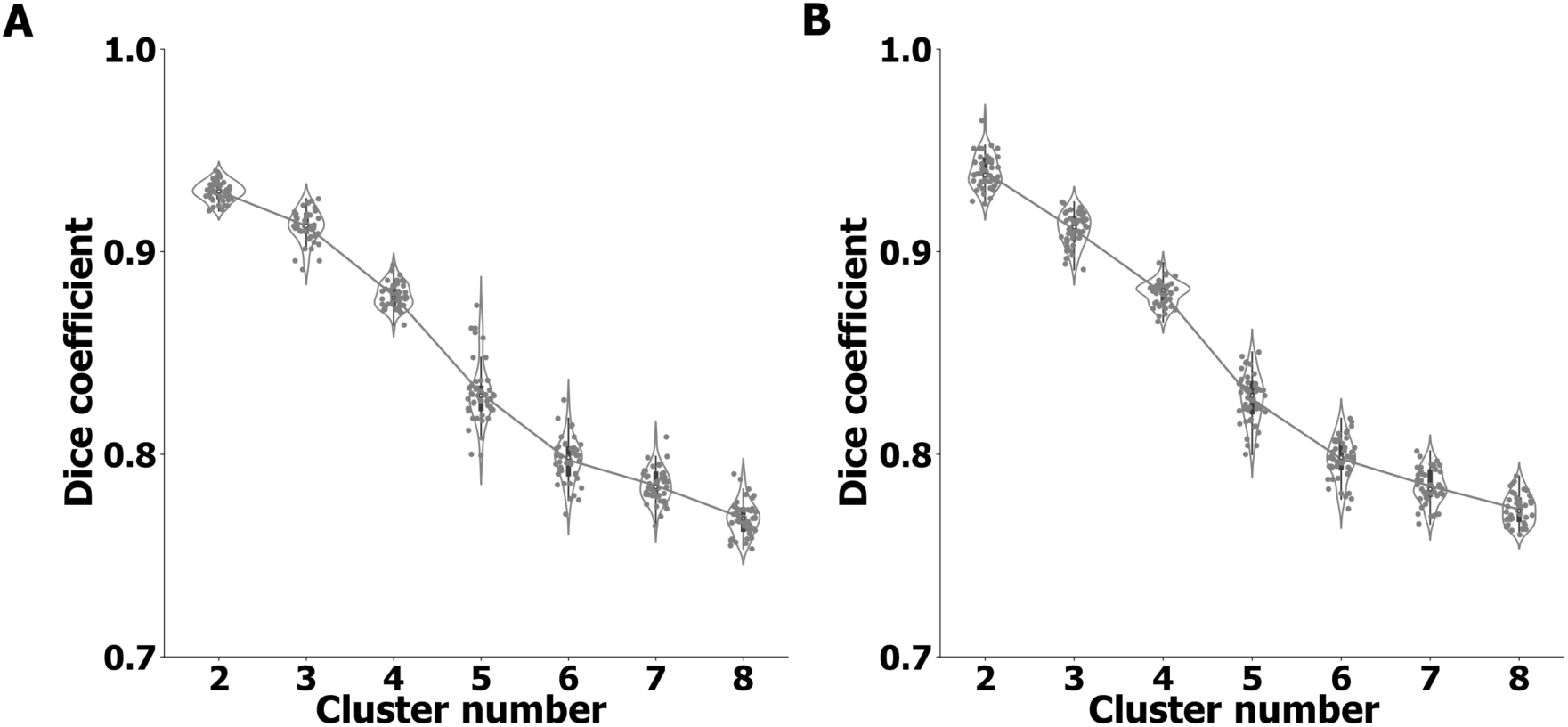
Stability analysis of identified clusters. The stability of the identified subtypes was evaluated using seed-based and subsample-based strategies, assessed by the Dice coefficient. To generate more than two subtypes, k-means clustering was applied to the predictive scores of CU individuals derived from classification models. **(A)** In the seed-based strategy, classification and clustering analyses were repeated across 10 trials, each initialized with a different random seed. **(B)** In the subsample-based strategy, 90% of participants were randomly subsampled in the classification step across 10 trials, followed by clustering analysis. For both methods, the Dice coefficient was used to quantify the consistency of clustering labels across trials, demonstrating the robustness of the identified subtypes.

**Figure S6.**
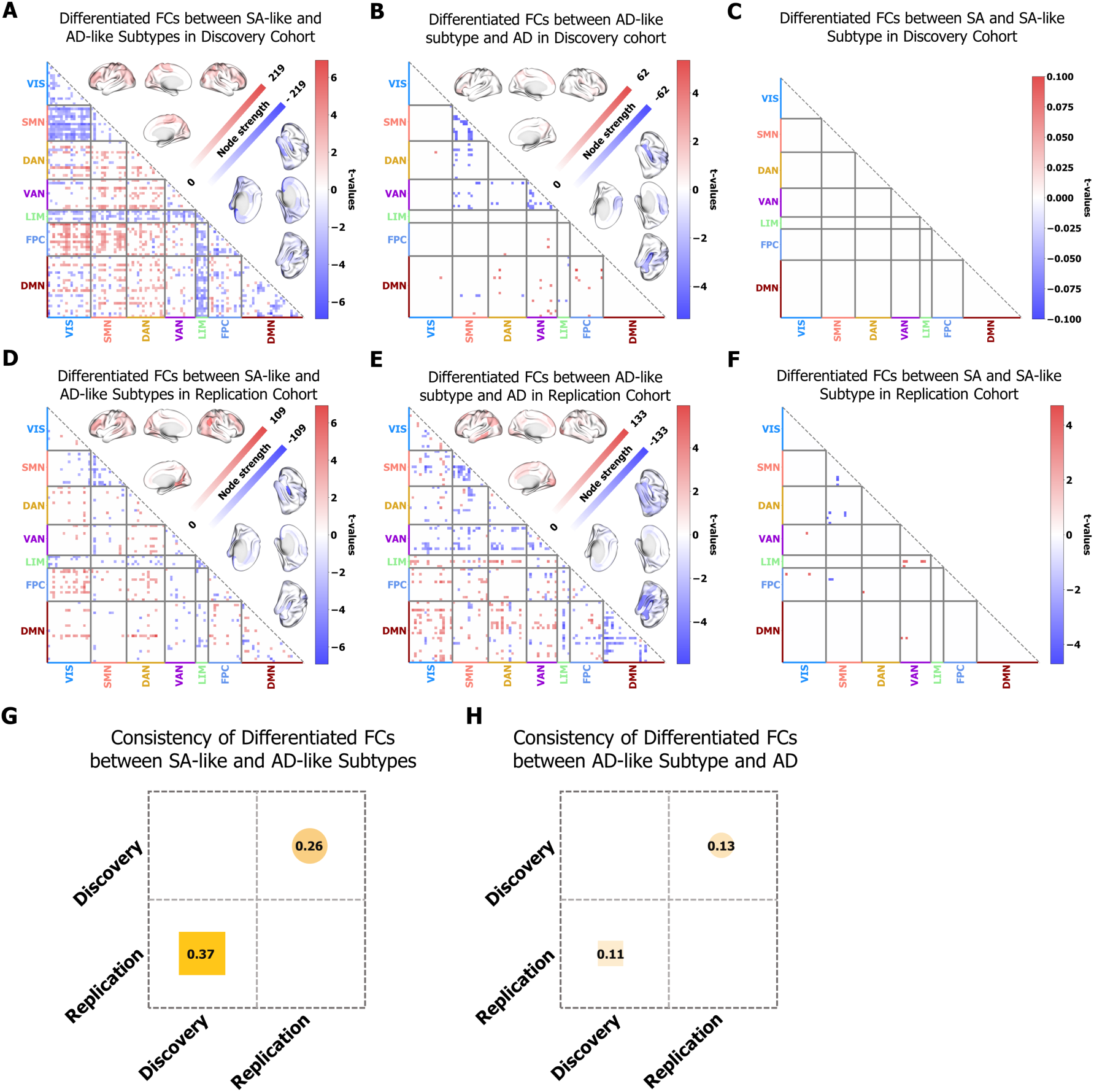
FCs differentiated across two subtypes, SA, and AD. We identified FC differences between SA-like and AD-like subtypes, AD-like subtype and AD, and SA-like subtype and SA in both discovery and replication cohorts, using two-sample t-tests while controlling for age and sex. The similarity of FC differences between the two cohorts was further assessed. **(A) (D)** Significantly (p_fdr_ ≤ 0.05) differentiated FCs between SA-like and AD-like subtypes in discovery and replication cohorts. **(B) (E)** Significantly (p_fdr_ ≤ 0.05) differentiated FCs between AD-like subtype and AD in discovery and replication cohorts. **(C) (F)** Significantly (p_fdr_ ≤ 0.05) differentiated FCs between SA and SA-like subtype in discovery and replication cohorts. **(G)** Consistency of FCs differences between SA-like and AD-like subtypes across discovery and replication cohorts, measured by Kendall correlation (circle box) and Dice coefficient (rectangle box). **(H)** Consistency of FCs differences between AD-like subtype and AD across discovery and replication cohorts. The significance of the consistency was confirmed by permutation test in 1000 trials. All consistent scores shown in **(G)**, **(H)** were significant (p ≤ 0.001). The consistency of FCs differentiating SA and SA-like subtypes was not measured, as no FCs significantly differentiated these groups in the discovery cohort.

**Figure S7.**
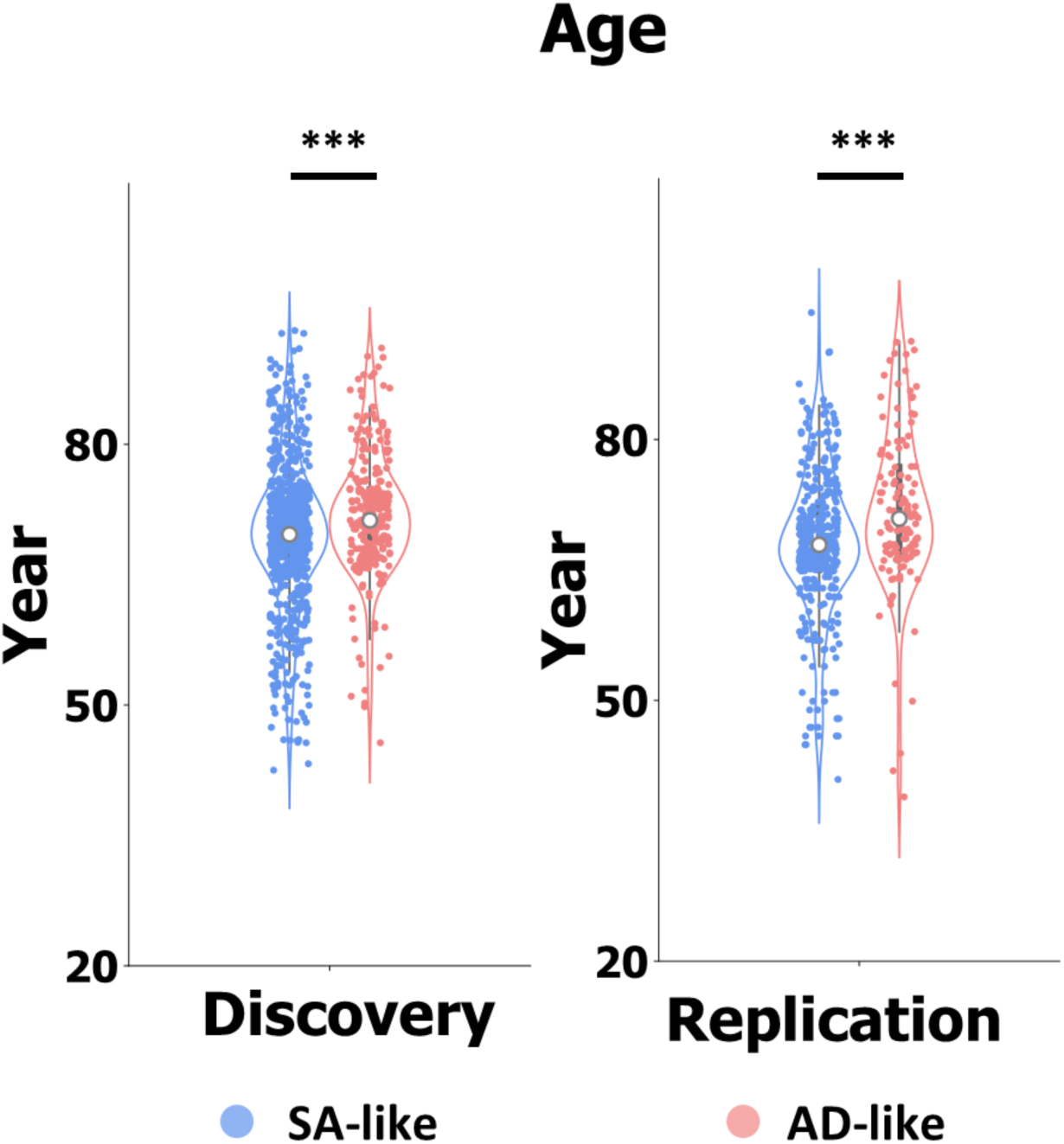
Distribution of age between SA-like and AD-like subtypes at baseline. We used Kruskal-Wallis test to compare the age of different subtypes. The significance was annotated by ***(p_fdr_ ≤ 0.001).

**Figure S8.**
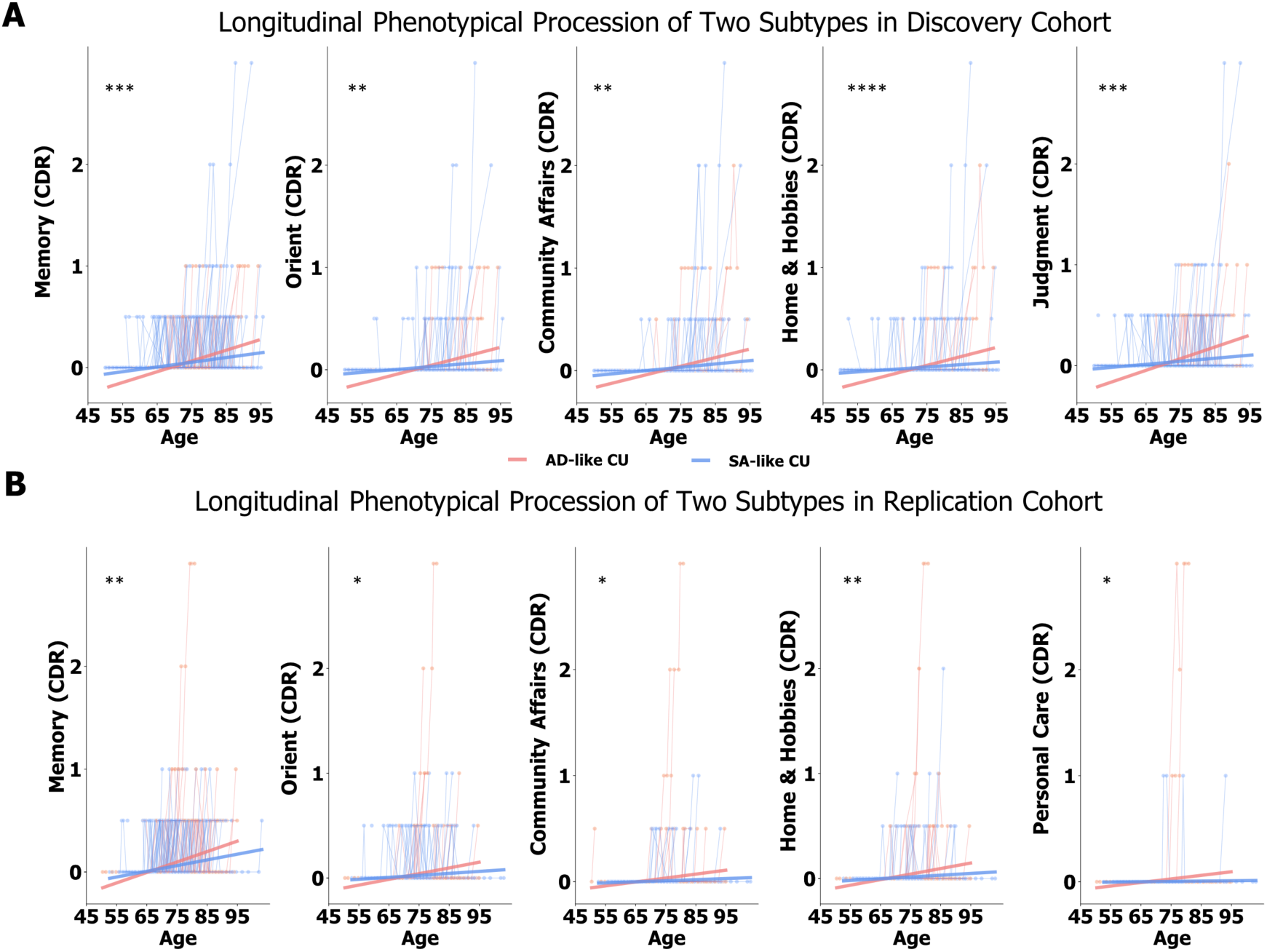
Longitudinal changes in item scores from CDR and FAQ differentiating SA-like and AD-like subtypes in discovery and replication cohorts. A linear mixed-effects model was used to assess the interaction between age and subtype on various item scores. Only items with significant interaction effects that explained variance in phenotypic measures are displayed. Significance levels are indicated as * (p_fdr_ ≤ 0.05), ** (p_fdr_ ≤ 0.01), *** (p_fdr_ ≤ 0.001), and **** (p_fdr_ ≤ 0.0001).

**Figure S9.**
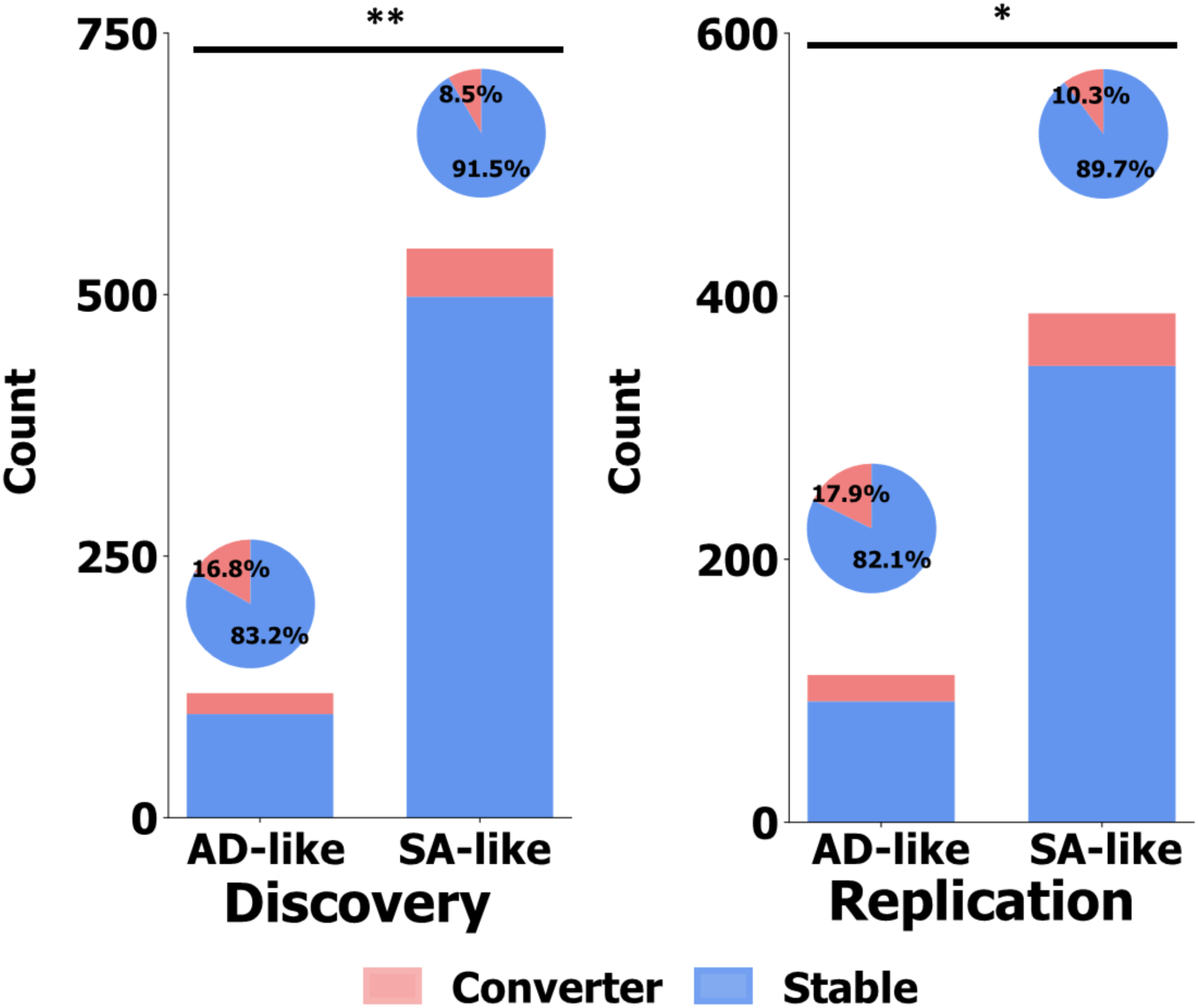
Conversion risk to MCI/AD differs between cognitive aging subtypes. For both SA-like and AD-like subtypes, bar plots display counts of individuals who remained CU (stable) or progressed to MCI/AD (converters) at follow-up, with pie charts showing proportions. A chi-square test was performed to assess the difference in conversion rates between the SA-like and AD-like subtypes (discovery cohort: χ^2^ = 6.7, p = 0.009; replication cohort: χ^2^ = 4.0, p = 0.04).

**Figure S10.**
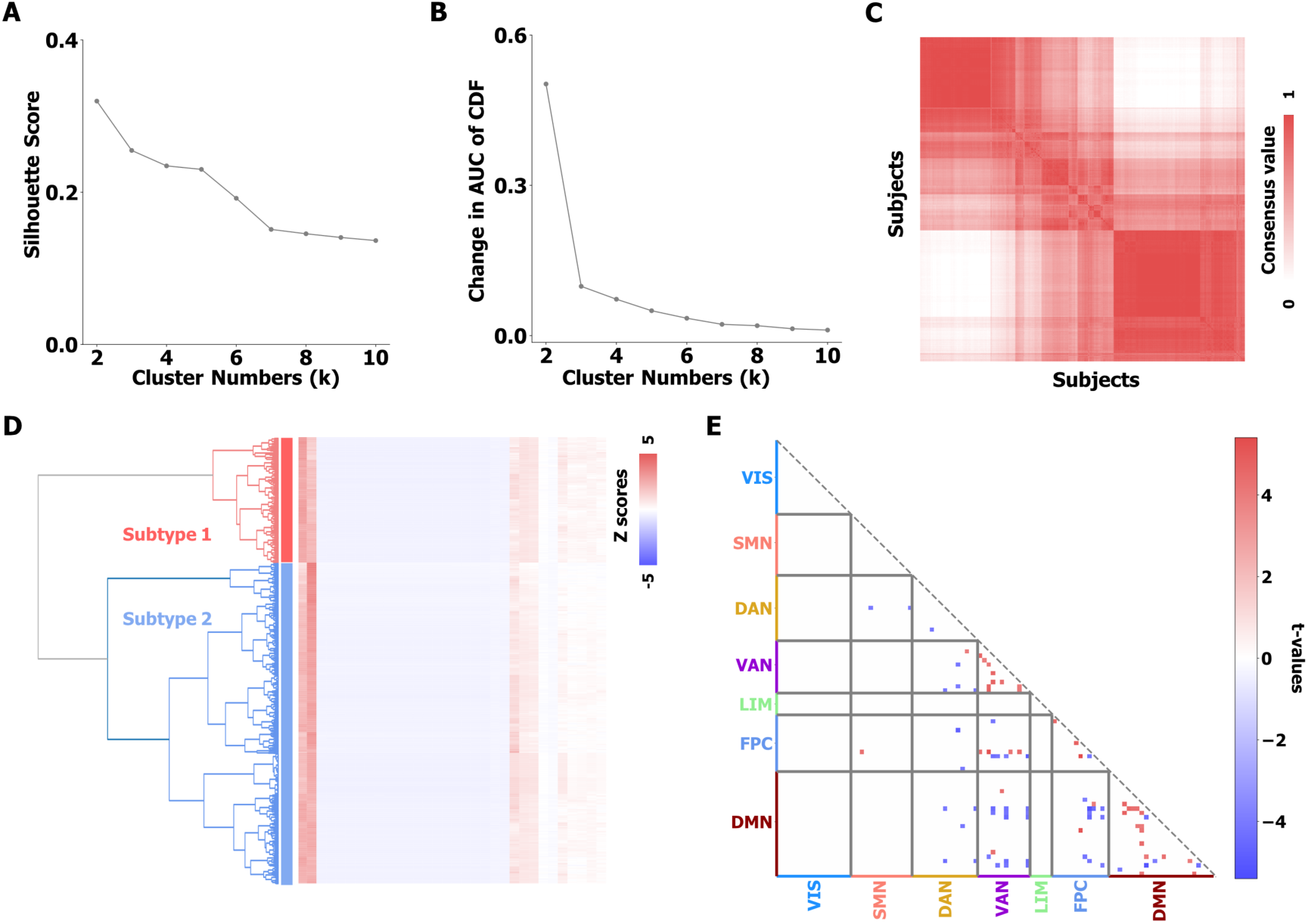
Clustering based on characteristic variables of the discovery cohort. We performed hierarchical clustering to identify subtypes based on characteristic variables and used consensus clustering to evaluate the stability of the subtyping. **(A)** Silhouette scores for different cluster numbers, reflecting clustering quality. Larger silhouette scores indicate better cluster quality. **(B)** Change in the area under the curve (AUC) of the cumulative distribution function (CDF) for consensus clustering across various cluster numbers, used to determine the most stable clustering solution. Larger changes in AUC of CDF indicate better stability of the identified clusters. **(C)** Ordered consensus matrix for K = 2, illustrating cluster stability. White indicates no consensus, while red indicates very high consensus. More elements centered along the diagonal indicate better stability. **(D)** Hierarchical clustering dendrogram, where the height of each linkage represents the distance between merged clusters. **(E)** FC differences between the two identified subtypes, measured by two-sample t-tests. Only FCs that survived false discovery rate correction are shown.

**Figure S11.**
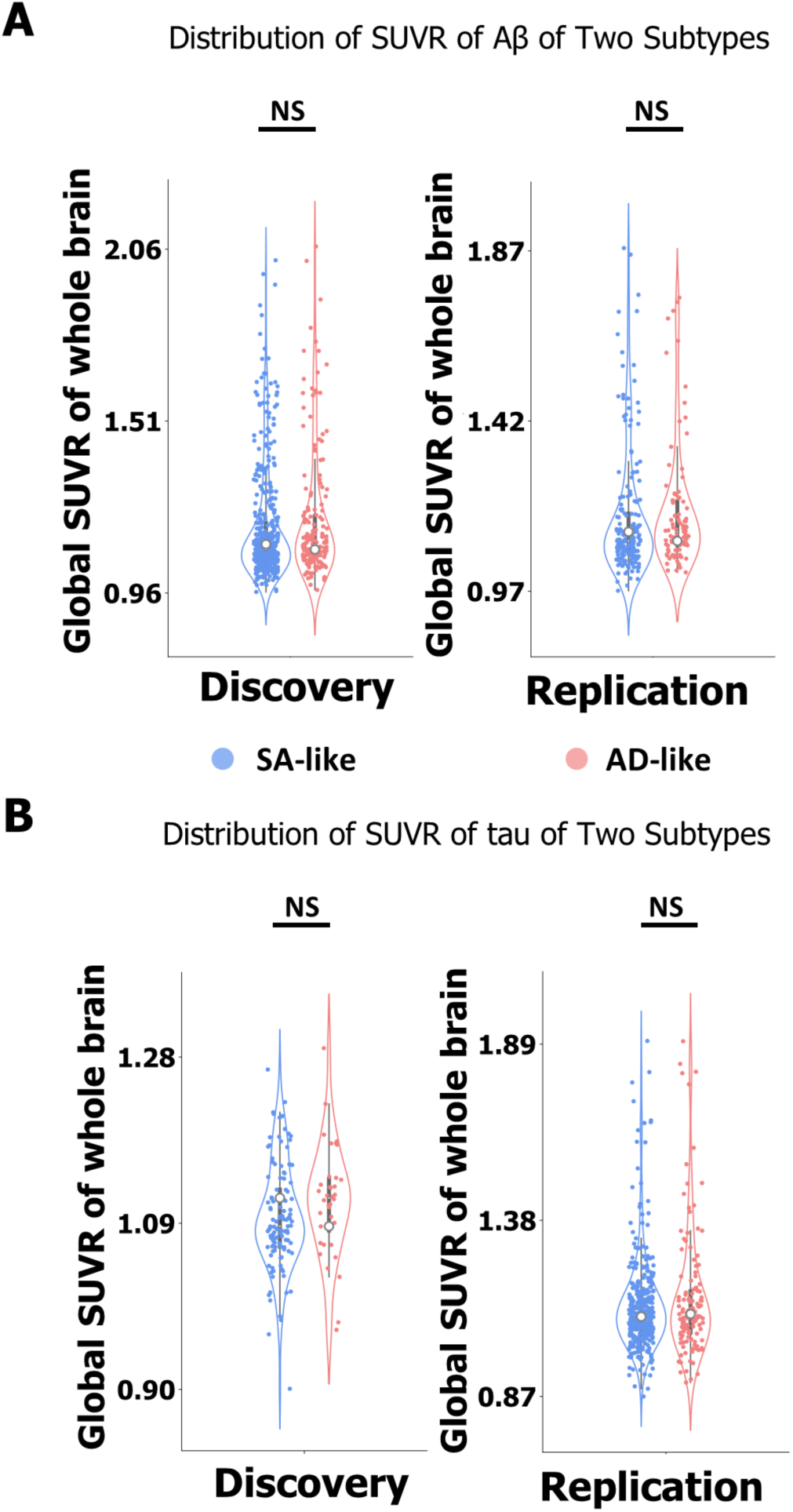
Distribution of neuropathological biomarkers between SA-like and AD-like subtypes at the baseline. We used Kruskal-Wallis test to compare the standardized uptake value ratio (SUVR) of Aβ and tau between identified subtypes. No significant results were found.

**Figure S12.**
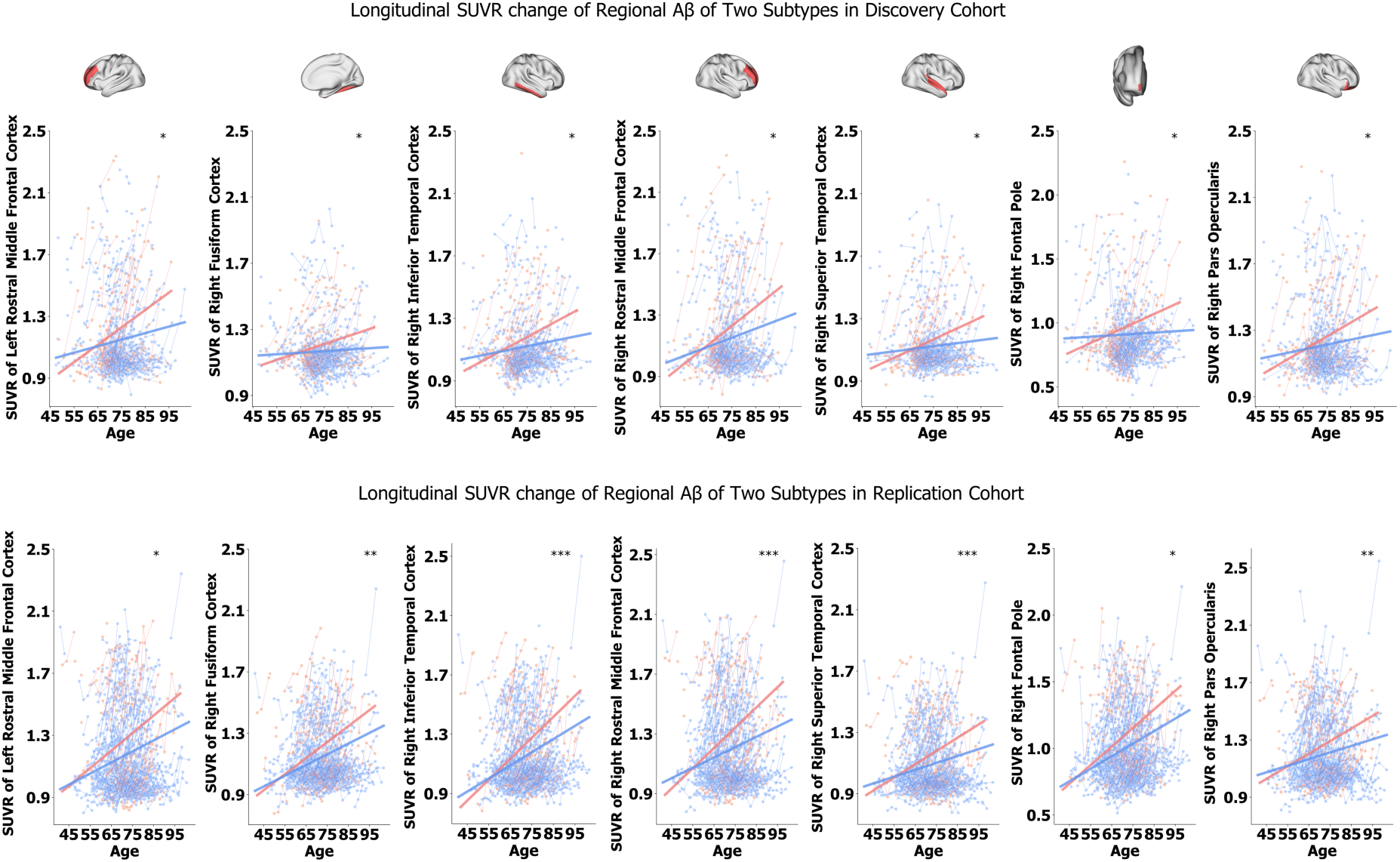
Longitudinal trajectories of regional Aβ expression between SA-like and AD-like subtypes in discovery and replication cohorts. A linear mixed-effects model was used to examine the time-by-subtype interaction effect on regional Aβ expression. The p-values from all brain regions in the discovery cohort were FDR-corrected, and only significant results were visualized with corresponding brain surface maps. Significance levels are indicated as * (p_fdr_ ≤ 0.05), ** (p_fdr_ ≤ 0.01), and *** (p_fdr_ ≤ 0.001). The longitudinal trends of the same regions were then plotted in the replication cohort to assess reproducibility.

**Figure S13.**
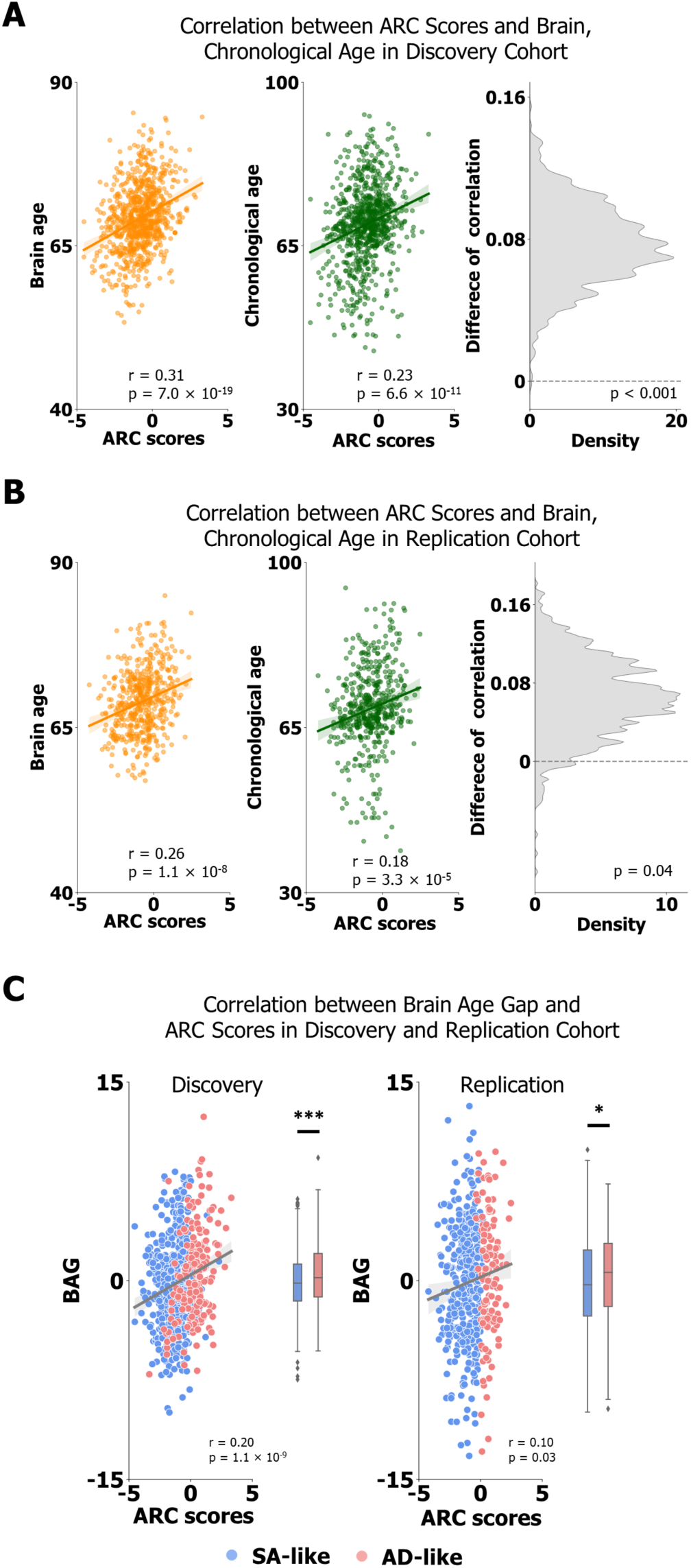
Association between the ARC signature and brain aging between two subtypes. We assessed the relationships between brain age, chronological age, and ARC signature scores (normalized weighted sum of ARC FCs). The difference between the correlation of brain age with ARC scores and that of chronological age with subtyping scores was evaluated using 1,000 bootstrapping trials. **(A)** Correlations between brain age, chronological age, and ARC scores in the discovery cohort. **(B)** Correlations between brain age, chronological age, and ARC scores in the replication cohort. **(C)** Association between brain age gap (BAG) and subtyping scores in the discovery cohort. BAG was calculated as the difference between brain age and chronological age. The difference of BAG between the two subtypes was confirmed by two-sample t-test, with significance levels denoted as * (p ≤ 0.05), and *** (p ≤ 0.001).

**Figure S14.**
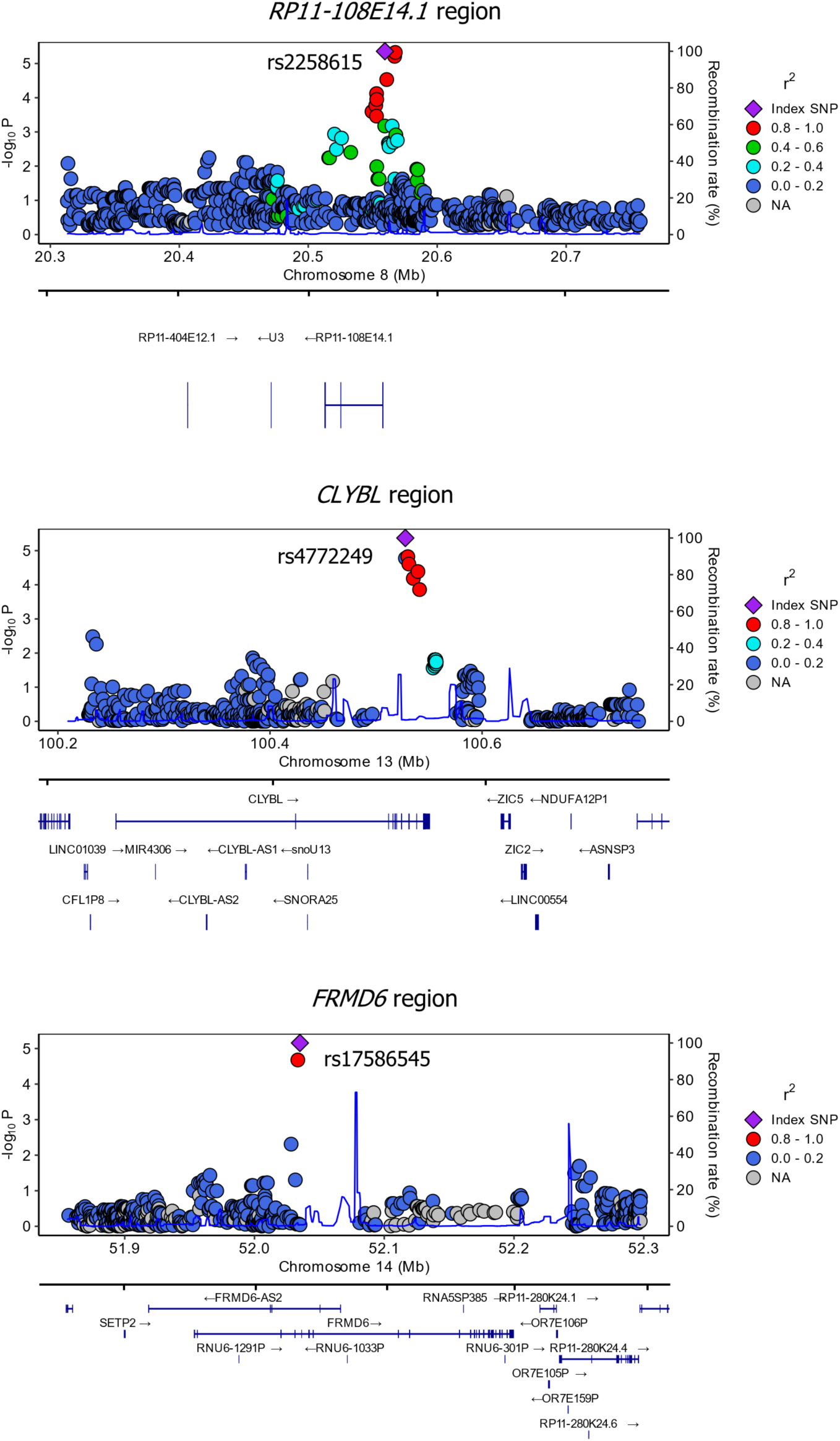
Locus plots for three loci differentiating between AD-like and SA-like subtypes. At each locus, genotyped SNPs are represented as purple boxes. Estimated recombination rates (derived from the 1000 Genomes European population (CEU)) are plotted to illustrate the local linkage disequilibrium (LD) structure. The color of the SNPs indicates their LD with the index SNP, based on pairwise r² values from the 1000 Genomes CEU individuals. Gene annotations were obtained from the Ensembl Genome Browser.

**Figure S15.**
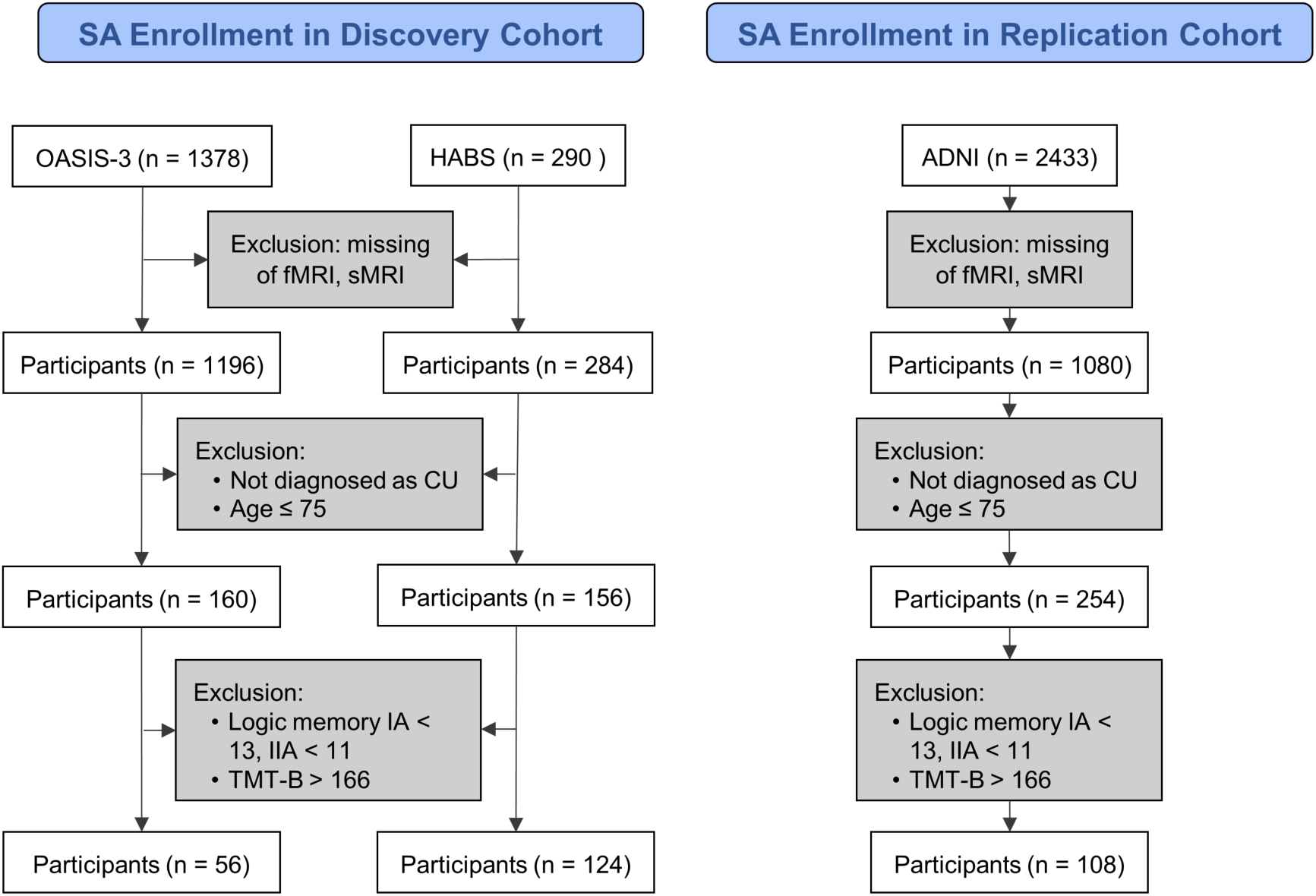
Flow diagram of SA inclusion in complementary analysis, using consistent memory measures across cohorts. The difference between this diagram and the one from Figure S1 is that WAIS-R Logical Memory was used instead of RAVLT to assess memory performance in the replication cohort, ensuring a consistent SA definition across cohorts.

**Figure S16.**
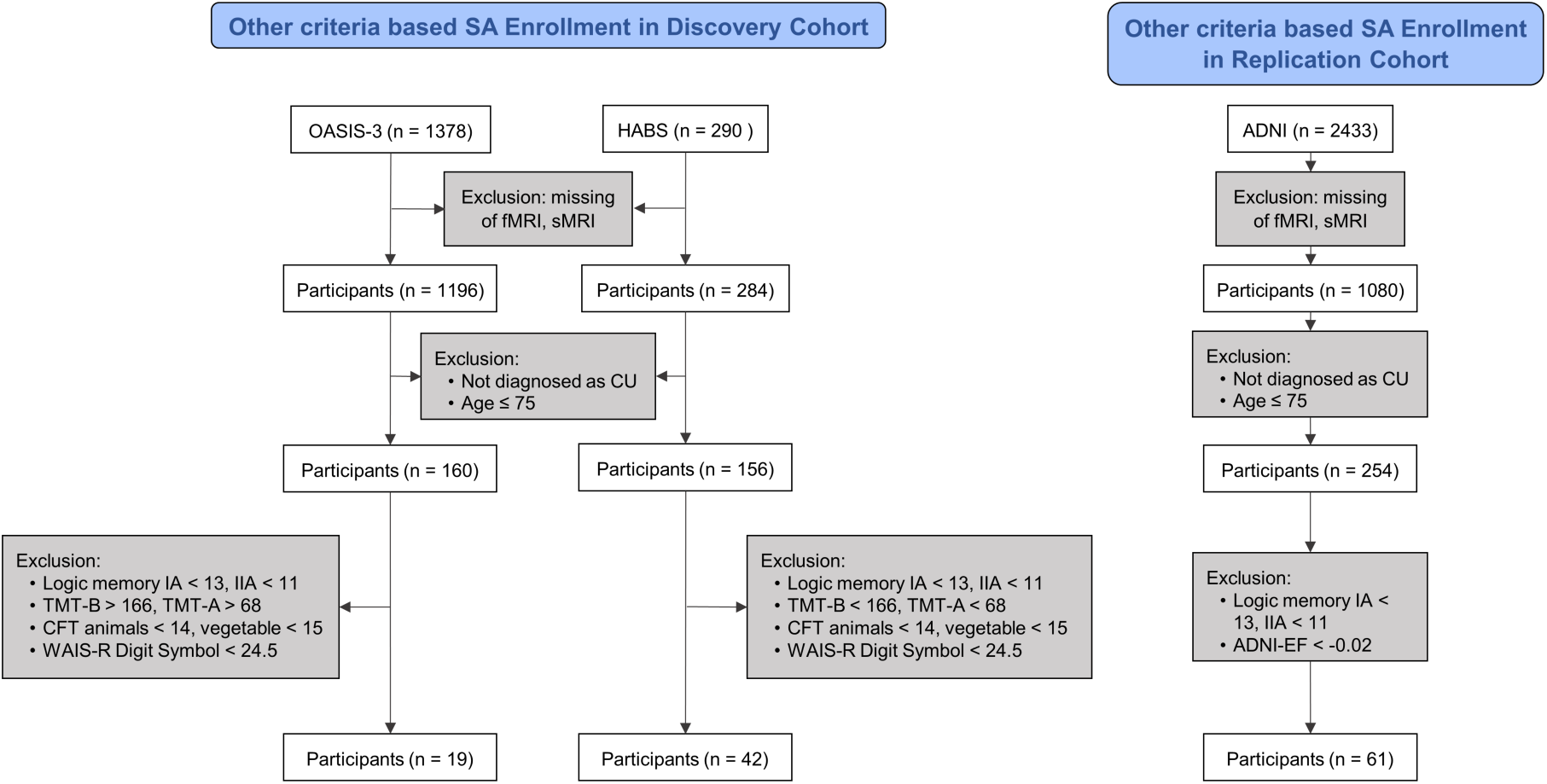
Flow diagram of SA inclusion in the second complementary analysis, using comprehensive cognitive tests measuring executive function to define SA.

**Figure S17.**
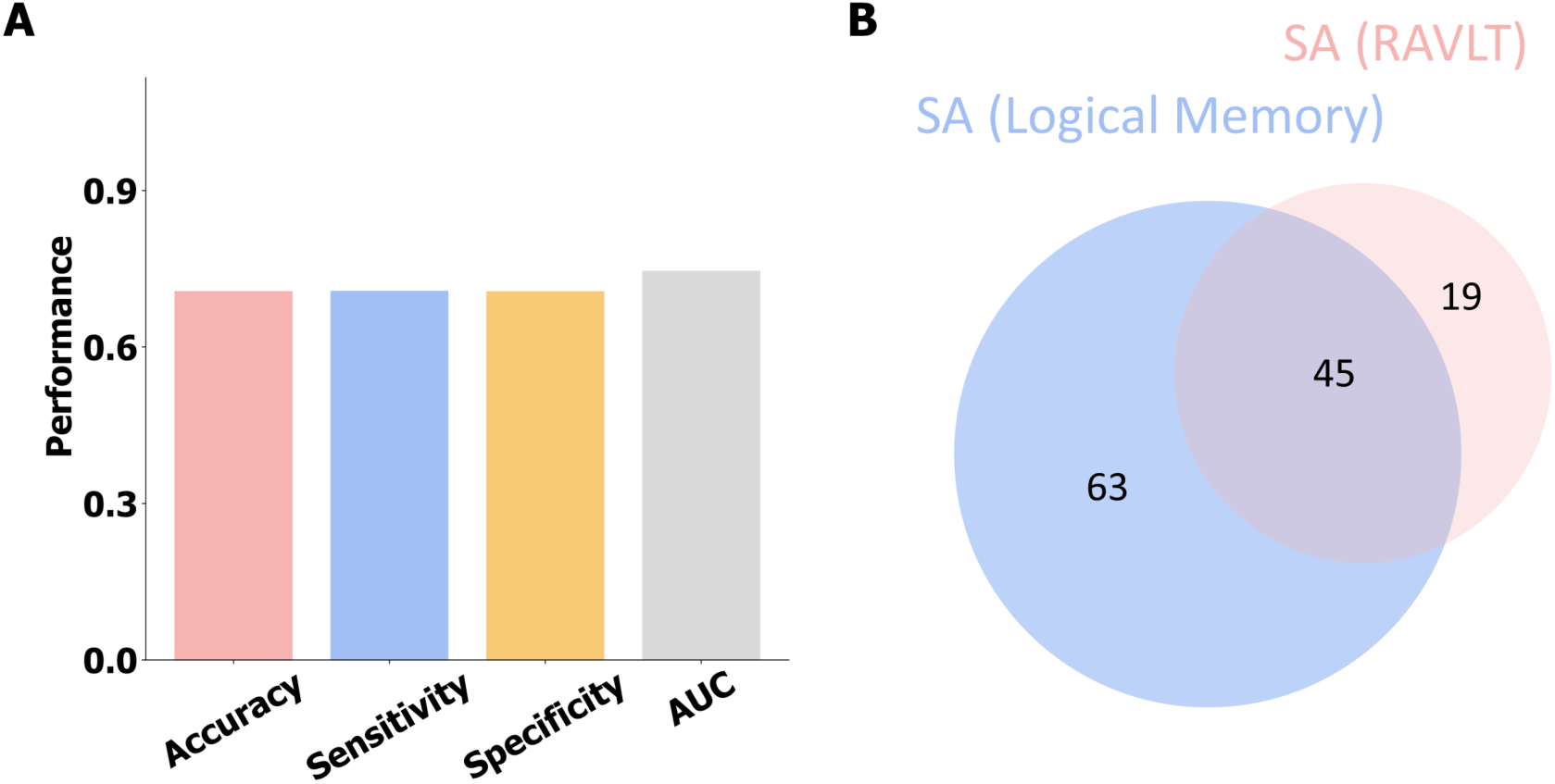
Classification between SA and AD defined using WAIS-R logic memory test. **(A)** Performance evaluation in the replication cohort using the classifier trained on the discovery cohort. **(B)** Venn diagram illustrating the overlap between SA identified by WAIS-R Logical Memory versus RAVLT criteria in the replication cohort.

**Figure S18.**
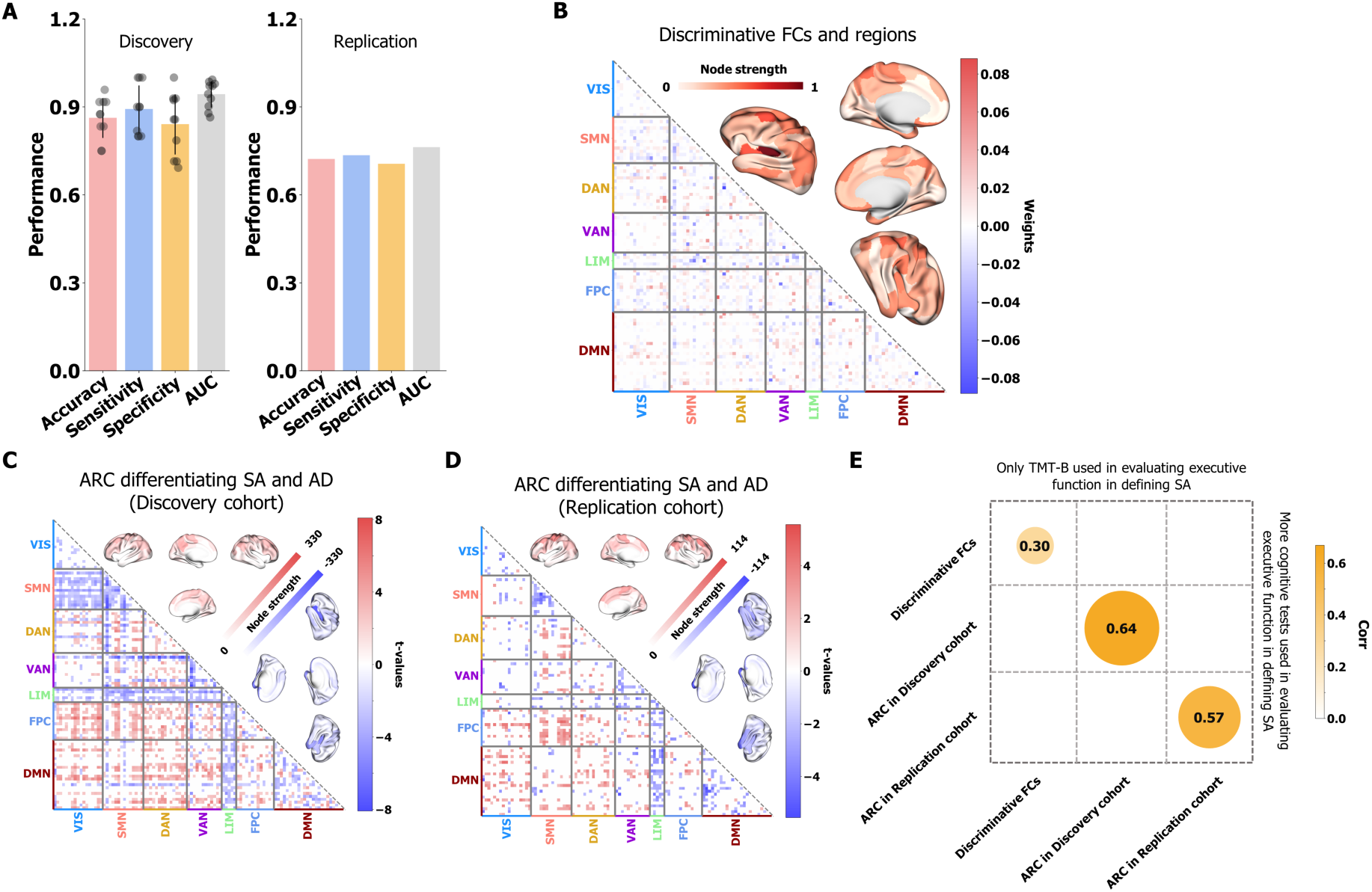
Identification and validation of the ARC distinguishing SA from AD, using more comprehensive executive function criteria for superager definition. **(A)** Classification performance in the discovery and replication cohorts. In the discovery cohort, tenfold cross-validation yielded an accuracy of 0.82 ± 0.07, sensitivity of 0.81 ± 0.06, specificity of 0.84 ± 0.12, and an area under the curve (AUC) of 0.89 ± 0.05. Applying the classifier to the replication cohort resulted in an accuracy of 0.76, sensitivity of 0.78, specificity of 0.74, and an AUC of 0.83. **(B)** Discriminative FC signature based on the average weights of the ten cross-validated classifiers. Node strength, defined as the sum of absolute FC classifier weights for each region, is visualized on the brain surface to highlight regions contributing most to classification. **(C), (D)** ARC signatures that significantly differentiated SA from AD as detected by two-sample t-tests (FDR-corrected p < 0.05) in the discovery and replication cohorts. Hypo-connections (stronger in SA) are colored red, while hyper-connections (stronger in AD) are colored blue. Node strength, derived from summed positive and negative t-values of discriminative FCs, is visualized on the brain surface. **(E)** High consistency of both discriminative FCs and ARC signatures between the two SA definitions, demonstrated by significant Kendall rank correlations (all p<0.0001).

**Table S1.** Demographic information of the discovery cohort. In the classification analysis, AD participants were matched based on an age cutoff (> 75 years). Differences in binary categorical variables between superagers and AD participants in the classification step were assessed using the Chi-squared test, while differences in ordinal categorical and continuous variables were evaluated using the Wilcoxon signed-rank test. All statistical comparisons were conducted as two-sided tests. BL (baseline) refers to the earliest enrollment date of the participant, which could be at the first visit or during subsequent sessions. The demographic information for CU (IV) excludes subjects who were classified as superagers after reaching the age of 75 and were used in classification step.

| Characteristic variables | Subtyping analysis |  |  |  |  |  |  |  |  |  |
| --- | --- | --- | --- | --- | --- | --- | --- | --- | --- | --- |
|  | Classification analysis |  |  |  |  |  |  |  |  |  |
|  | SA |  | AD (SA age matched) |  | Statistic values |  | CU (BL) |  | AD (BL, excluding SA age matched) |  |
| | n/median | %/IQR | n/median | %/IQR | $\chi^2/Z$ | p | n/median | %/IQR | n/median | %/IQR |
| Sex (Male) | 73 | 41 | 21 | 20 | 14 | 0.0003 | 310 | 38 | 32 | 30 |
| Sex (Female) | 107 | 59 | 89 | 80 |  |  | 508 | 62 | 73 | 70 |
| Age | 80 | [77, 82] | 80 | [77, 83] | -0.28 | 0.78 | 69 | [66, 75] | 69 | [66, 72] |
| Education years | 16 | [14, 18] | 16 | [14, 18] | 1.54 | 0.12 | 16 | [14, 18] | 16 | [14, 18] |
| Race (White) | 177 | 98 | 94 | 85 | 16.5 | $5.0 \times 10^{-6}$ | 705 | 86 | 77 | 73 |
| Race (Black or African American) | 2 | 1 | 14 | 13 |  |  | 82 | 10 | 8 | 8 |
| Race (Asian) | 0 | 0 | 0 | 0 |  |  | 8 | 1 | 2 | 2 |
| Race (Unkown) | 1 | 1 | 2 | 2 |  |  | 23 | 3 | 17 | 17 |

**Table S2.** Demographic information of the replication cohort. In the classification analysis, AD participants were matched based on an age cutoff (> 75 years). Differences in binary categorical variables between superagers and AD participants in the classification step were assessed using the Chi-squared test, while differences in ordinal categorical and continuous variables were evaluated using the Wilcoxon signed-rank test. All statistical comparisons were conducted as two-sided tests. BL (baseline) refers to the earliest enrollment date of the participant, which could be at the first visit or during subsequent sessions. The demographic information for CU (IV) excludes subjects who were classified as superagers after reaching the age of 75 and were used in classification step.

| Characteristic variables | Subtyping analysis |  |  |  |  |  |  |  |  |  |
| --- | --- | --- | --- | --- | --- | --- | --- | --- | --- | --- |
|  | Classification analysis |  |  |  |  |  |  |  |  |  |
|  | SA |  | AD (SA age matched) |  | Statistic values |  | CU (BL) |  | AD (BL, excluding SA age matched) |  |
| | n/median | %/IQR | n/median | %/IQR | $\chi^2/Z$ | p | n/median | %/IQR | n/median | %/IQR |
| Sex (Male) | 22 | 36 | 47 | 59 | 8 | 0.005 | 202 | 41 | 25 | 50 |
| Sex (Female) | 42 | 64 | 32 | 41 |  |  | 295 | 59 | 25 | 50 |
| Age | 79 | [77, 83] | 80 | [77, 85] | -1.1 | 0.26 | 69 | [65, 74] | 70 | [64, 72] |
| Education years | 18 | [16, 18] | 16 | [14, 17] | 4.2 | $2.7 \times 10^{-4}$ | 16 | [15, 18] | 16 | [14, 18] |
| Race (White) | 57 | 89 | 71 | 90 | 0 | 1 | 394 | 79 | 43 | 86 |
| Race (Black or African American) | 6 | 9 | 3 | 4 |  |  | 65 | 13 | 6 | 12 |
| Race (Asian) | 1 | 2 | 4 | 5 |  |  | 20 | 4 | 1 | 2 |
| Race (Unkown) | 0 | 0 | 1 | 1 |  |  | 18 | 4 | 0 | 0 |

**Table S3.** Demographic information of the identified subtypes in the discovery and replication cohorts. Categorical variables such as sex and race between superager-like and AD-like cognitive unimpaired subtypes were compared using the Chi-squared test, while differences in ordinal categorical and continuous variables between two subtypes were evaluated using Kruskal-Wallis test. All statistical comparisons were conducted as two-sided tests. FDR correction was applied to p values for all variables, including cognitive measures listed in Tables S4 and S5, separately for the discovery and replication cohorts.

| Variables | Discovery cohort |  |  |  |  |  | Replication cohort |  |  |  |  |  |
| --- | --- | --- | --- | --- | --- | --- | --- | --- | --- | --- | --- | --- |
|  | SA-like CU<br>(n = 607) |  | AD-like CU<br>(n = 211) |  | Statistic<br>values |  | SA-like CU<br>(n = 370) |  | AD-like CU<br>(n = 127) |  | Statistic<br>values |  |
| | n/median | %/IQR | n/median | %/IQR | $\chi^2/Z$ | p<sub>fdr</sub> | n/median | %/IQR | n/median | %/IQR | $\chi^2/Z$ | p<sub>fdr</sub> |
| Age | 69 | [65, 74] | 71 | [67, 75] | 14.3 | **0.0002** | 68 | [65, 73] | 71 | [67, 77] | 19.6 | **0.0002** |
| Sex (Male) | 61 | 10 | 22 | 10 | 0 | 1 | 144 | 39 | 58 | 46 | 1.52 | 0.45 |
| Sex (Female) | 546 | 90 | 189 | 90 |  |  | 226 | 61 | 69 | 54 |  |  |
| Education years | 16 | [14, 18] | 16 | [14, 18] | 0.01 | 0.91 | 16 | [16, 18] | 17 | [15, 18] | 0.05 | 0.95 |
| Race (White) | 528 | 87 | 178 | 84 | 0.70 | 0.50 | 222 | 60 | 75 | 59 | 0.04 | 0.95 |
| Race (Others) | 79 | 13 | 33 | 16 |  |  | 148 | 40 | 52 | 41 |  |  |

**Table S4.** Statistical comparison of the characteristic variables between two identified subtypes at baseline from discovery cohort. Differences in ordinal categorical and continuous variables between the two subtypes were evaluated using Kruskal-Wallis test. All statistical comparisons were conducted as two-sided tests. FDR was used to correct p values of comparison of all variables, including the demographic variables in Table S3. NaN result means the variables of two subtypes were constant and same.

| Characteristic variables | Statistic values |  |  |  |
| --- | --- | --- | --- | --- |
| | $\chi^2$ | p<sub>fdr</sub> | n<sub>1</sub> (SA-like CU) | n<sub>2</sub> (AD-like CU) |
| Community Affairs (CDR) | 0.24 | 0.62 | 356 | 87 |
| Home & Hobbies (CDR) | 0.74 | 0.39 | 356 | 87 |
| Personal Care (CDR) | NaN | NaN | 356 | 87 |
| Judgment (CDR) | 0.13 | 0.72 | 356 | 87 |
| Memory (CDR) | 1.13 | 0.29 | 514 | 145 |
| Orientation (CDR) | 0.24 | 0.62 | 356 | 87 |
| SOB (CDR) | 0.04 | 0.83 | 514 | 145 |
| TMT-A (NAB) | 0.52 | 0.47 | 372 | 120 |
| TMT-B (NAB) | 1.05 | 0.31 | 371 | 120 |
| RAVLT-Immediate (NAB) | NaN | NaN | 0 | 0 |
| RAVLT-Learning (NAB) | NaN | NaN | 0 | 0 |
| RAVLT-Perc-Forgetting (NAB) | NaN | NaN | 0 | 0 |
| Logical Memory Delayed (NAB) | 1.34 | 0.25 | 372 | 120 |
| Logical Memory Immediate (NAB) | 2.56 | 0.11 | 372 | 120 |
| CFT-Animals (NAB) | 0.74 | 0.39 | 372 | 120 |
| CFT-Vegetable (NAB) | 0.05 | 0.82 | 372 | 120 |
| Digit Span-Forward (NAB) | 0.49 | 0.48 | 214 | 62 |
| Digit Span-Backward (NAB) | 2.79 | 0.09 | 214 | 62 |
| Boston Naming Test (NAB) | 0.22 | 0.64 | 214 | 62 |
| Paying Bills (FAQ) | 0.86 | 0.35 | 313 | 67 |
| Taxes and Business Affairs (FAQ) | 0.11 | 0.74 | 300 | 68 |
| Shopping Alone (FAQ) | 0.22 | 0.64 | 326 | 72 |
| Games and Hobbies (FAQ) | 0.22 | 0.64 | 302 | 65 |
| Using Stove (FAQ) | 0.22 | 0.64 | 328 | 72 |
| Preparing a Balanced Meal (FAQ) | NaN | Nan | 309 | 68 |
| Current Events (FAQ) | 0.44 | 0.51 | 327 | 72 |
| Paying Attention (FAQ) | 1.40 | 0.24 | 329 | 72 |
| Remembering Dates (FAQ) | 0.44 | 0.51 | 329 | 72 |
| Traveling and Driving (FAQ) | 0.07 | 0.79 | 327 | 72 |

**Table S5.**
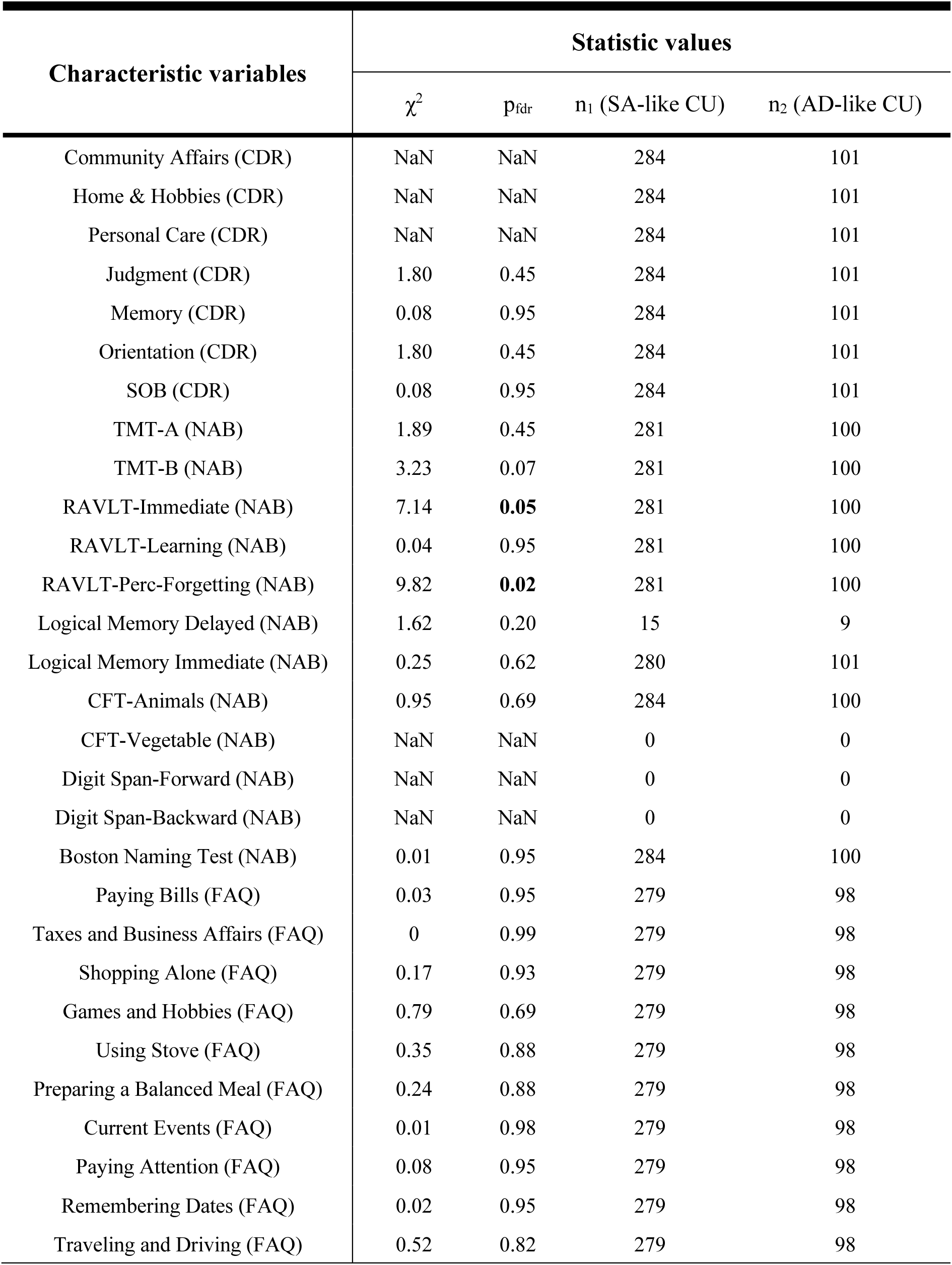
Statistical comparison of the characteristic variables between two identified subtypes at baseline from replication cohort. Categorical variables such as sex and race between superager-like and AD-like cognitive unimpaired subtypes were compared using the Chi-squared test, while differences in ordinal categorical and continuous variables between two subtypes were evaluated using Kruskal-Wallis test. All statistical comparisons were conducted as two-sided tests. FDR were used to correct p values of comparison of all variables.

| Characteristic variables | Statistic values |  |  |  |
| --- | --- | --- | --- | --- |
| | $\chi^2$ | p<sub>fdr</sub> | n<sub>1</sub> (SA-like CU) | n<sub>2</sub> (AD-like CU) |
| Community Affairs (CDR) | NaN | NaN | 284 | 101 |
| Home & Hobbies (CDR) | NaN | NaN | 284 | 101 |
| Personal Care (CDR) | NaN | NaN | 284 | 101 |
| Judgment (CDR) | 1.80 | 0.45 | 284 | 101 |
| Memory (CDR) | 0.08 | 0.95 | 284 | 101 |
| Orientation (CDR) | 1.80 | 0.45 | 284 | 101 |
| SOB (CDR) | 0.08 | 0.95 | 284 | 101 |
| TMT-A (NAB) | 1.89 | 0.45 | 281 | 100 |
| TMT-B (NAB) | 3.23 | 0.07 | 281 | 100 |
| RAVLT-Immediate (NAB) | 7.14 | **0.05** | 281 | 100 |
| RAVLT-Learning (NAB) | 0.04 | 0.95 | 281 | 100 |
| RAVLT-Perc-Forgetting (NAB) | 9.82 | **0.02** | 281 | 100 |
| Logical Memory Delayed (NAB) | 1.62 | 0.20 | 15 | 9 |
| Logical Memory Immediate (NAB) | 0.25 | 0.62 | 280 | 101 |
| CFT-Animals (NAB) | 0.95 | 0.69 | 284 | 100 |
| CFT-Vegetable (NAB) | NaN | NaN | 0 | 0 |
| Digit Span-Forward (NAB) | NaN | NaN | 0 | 0 |
| Digit Span-Backward (NAB) | NaN | NaN | 0 | 0 |
| Boston Naming Test (NAB) | 0.01 | 0.95 | 284 | 100 |
| Paying Bills (FAQ) | 0.03 | 0.95 | 279 | 98 |
| Taxes and Business Affairs (FAQ) | 0 | 0.99 | 279 | 98 |
| Shopping Alone (FAQ) | 0.17 | 0.93 | 279 | 98 |
| Games and Hobbies (FAQ) | 0.79 | 0.69 | 279 | 98 |
| Using Stove (FAQ) | 0.35 | 0.88 | 279 | 98 |
| Preparing a Balanced Meal (FAQ) | 0.24 | 0.88 | 279 | 98 |
| Current Events (FAQ) | 0.01 | 0.98 | 279 | 98 |
| Paying Attention (FAQ) | 0.08 | 0.95 | 279 | 98 |
| Remembering Dates (FAQ) | 0.02 | 0.95 | 279 | 98 |
| Traveling and Driving (FAQ) | 0.52 | 0.82 | 279 | 98 |

**Table S6.** Statistical comparison of the characteristic variables between two identified subtypes defined using characteristic variables at baseline from discovery cohort. Categorical variables such as sex and race between two subtypes were compared using the Chi-squared test, while differences in ordinal categorical and continuous variables between the two subtypes were evaluated using Kruskal-Wallis test. All statistical comparisons were conducted as two-sided tests. FDR were used to correct p values of comparison of all variables. The overall sample size of CU in this analysis is larger than that in Tables S3 and S4 because individuals who completed only cognitive assessments without MRI scanning were included.

| Characteristic variables | Statistic values |  |  |  |
| --- | --- | --- | --- | --- |
| | $\chi^2/f$ | p<sub>fdr</sub> | n<sub>1</sub> (subtype1) | n<sub>2</sub> (subtype2) |
| Age | 12.6 | **0.001** | 339 | 867 |
| Sex | 0.09 | 0.77 | 339 | 867 |
| Race | NaN | NaN | 339 | 867 |
| Education years | 32.6 | **$4.4 \times 10^{-8}$** | 339 | 867 |
| Community Affairs (CDR) | 0.22 | 0.69 | 79 | 364 |
| Home & Hobbies (CDR) | 0.65 | 0.66 | 79 | 364 |
| Personal Care (CDR) | NaN | NaN | 79 | 364 |
| Judgment (CDR) | 1.54 | 0.40 | 79 | 364 |
| Memory (CDR) | 0.35 | 0.69 | 317 | 730 |
| Orientation (CDR) | 0.22 | 0.69 | 79 | 364 |
| SOB (CDR) | 4.66 | 0.07 | 317 | 730 |
| TMT-A (NAB) | 244 | **$8.1 \times 10^{-54}$** | 310 | 570 |
| TMT-B (NAB) | 552 | **$1.4 \times 10^{-123}$** | 310 | 569 |
| Logical Memory Delayed (NAB) | 89.5 | **$2.3 \times 10^{-20}$** | 310 | 569 |
| Logical Memory Immediate (NAB) | 84.5 | **$2.4 \times 10^{-19}$** | 310 | 570 |
| CFT-Animals (NAB) | 106 | **$6.4 \times 10^{-24}$** | 310 | 569 |
| CFT-Vegetable (NAB) | 55.2 | **$5.6 \times 10^{-13}$** | 310 | 569 |
| Digit Span-Forward (NAB) | 15.4 | **0.0003** | 72 | 204 |
| Digit Span-Backward (NAB) | 6.90 | **0.02** | 72 | 204 |
| Boston Naming Test (NAB) | 4.19 | 0.09 | 72 | 204 |
| Paying Bills (FAQ) | 0.11 | 0.77 | 71 | 309 |
| Taxes and Business Affairs (FAQ) | 0.91 | 0.57 | 68 | 300 |
| Shopping Alone (FAQ) | 0.22 | 0.69 | 71 | 327 |
| Games and Hobbies (FAQ) | 0.24 | 0.69 | 70 | 297 |
| Using Stove (FAQ) | 0.22 | 0.69 | 73 | 327 |
| Preparing a Balanced Meal (FAQ) | NaN | NaN | 66 | 311 |
| Current Events (FAQ) | 0.46 | 0.69 | 74 | 325 |
| Paying Attention (FAQ) | 1.33 | 0.44 | 74 | 327 |
| Remembering Dates (FAQ) | 0.49 | 0.69 | 74 | 327 |
| Traveling and Driving (FAQ) | 1.62 | 0.40 | 74 | 325 |

